# Seeing in the Dark: Intelligent Fourier Light Field Imaging for Bioluminescence Microscopy

**DOI:** 10.1101/2025.10.13.680966

**Authors:** Luis Felipe Morales Curiel, Aaron Kreis, Michael Krieg

**Affiliations:** Institut de Ciències Fotòniques, Castelldefels, The Barcelona Institute of Science and Technology, Barcelona, Spain

## Abstract

Bioluminescence microscopy offers a uniquely non-invasive window into cellular dynamics, yet its use has traditionally been limited by the intrinsically low brightness of luciferases. The poor photon budget forces long exposures, preventing faithful visualization of rapid physiological processes, especially in three dimensions. To overcome this barrier, we developed a Fourier light field microscope coupled with deep-learning–based reconstruction that achieves sub-second volumetric bioluminescence imaging, enabling direct recovery of 3D volumes from photon-limited measurements while suppressing reconstruction noise. This approach substantially mitigates the speed–resolution trade-off of conventional light field methods and bypasses the need for slow classical deconvolution priors on the depth-dependent point-spread function. We demonstrate its power by performing near real-time 3D calcium imaging in muscles and neurons of freely moving *Caenorhabditis elegans*, and by quantifying cell dynamics within stem-cell–derived spheroids using fluorescently labeled nuclei. Together, these results establish our framework as a practical tool for dynamic, volumetric studies of living systems.

## Introduction

Seeing biological processes unfold deep inside living tissues, ideally label-free with no added light and minimal perturbation, is a longstanding goal in bioimaging. Autofluorescence of biological tissues and living cells has been used as a contrast agent,^1^ but often requires ultraviolet excitation and prohibitively high illumination intensities due to the poor quantum yield of endogenous fluorophores.^2^ Since the cloning of the green fluorescent protein (GFP), specific labels can be attached to individual biomolecules which afforded swift recording of biological dynamics in three dimensions in a fluorescent confocal or light sheet microscope. However, the need to introduce tags onto endogenous molecules or use fixed tissues, pose significant barriers to truly non-invasive imaging. An additional complication is the dependence on external illumination, which can generate spurious background signals from autofluorescence, introduce phototoxic effects,^3^ and interfere with the intrinsic light sensitivity of biological specimens ranging from whole animals to subcellular organelles.

Bioluminescence offers an elegant solution through which proteins enzymatically produce light directly within the organism, without the need for an external excitation source.^4^ However, the catalytic turnover of such enzymes, hence photon production, is slow such that bioluminescence imaging is traditionally constrained by the long exposure times required to collect enough photons with an acceptable signal-to-noise ratio.^5^ Recently, we have shown that simple modification of the imaging protocols and technology and by combining the undersampled noisy images with content-aware restoration pipelines,^6^ afforded a strong reduction exposure times down to 50 ms.^5^ This balance of exposure and signal/noise ratio, accelerated by deep learning tech, was key to unlocking the second generation widefield bioluminescence microscopy. Intriguingly, the use of single-exposure light field imaging naturally complements bioluminescence by minimizing acquisition time and enabling 3D bioluminescence microscopy of subsecond calcium dynamics in freely moving animals.

Light field microscopy (LFM) improves this by capturing spatial and angular information using microlens arrays, enabling 3D reconstruction from a single exposure.^5,7,8^ LFM is particularly well suited for photon-starved images, as a single exposure is able to record all the directional and spatial information of the imaged scene. This approach pushes the boundaries of what can be observed in living systems using excitation-free imaging modalities. Unfortunately, scanless 3-D imaging with LFM comes with a cost; lateral spatial resolution must be sacrificed to record angular information. Therefore, a trade-off between lateral and axial resolution is made, where the loss of lateral resolution is proportional to the number of discrete angular samples acquired.^9^ This is because in standard LFM, spatial and angular information is directly mixed in the image plane, meaning that each microlens captures a different spatial region at multiple angles. This inherently reduces spatial resolution because pixels must be shared between these two types of information. A solution to this is Fourier light field microscopy (FLFM), in which the microlens array samples the light field in Fourier space,^10,11^ meaning the angular spectrum (light ray direction) is now encoded within each microlens’s aperture while sampling spatial details at higher frequencies. Although FLFM, like conventional LFM, sacrifices lateral resolution for single-shot volumetric acquisition because each microlens samples only a fraction of the objective pupil, this Fourier-space configuration offers important advantages. Spatial resolution remains more uniform across the imaging volume, and reconstruction artifacts at the native focal plane are largely eliminated because the recorded angular information remains non-redundant. This preserves higher lateral spatial sampling compared to conventional LFM while extending depth of field (DoF).

To visualize three-dimensional biological structures using LFM, raw image data must be computationally reconstructed into focused image planes. This involves disentangling spatial and angular light information captured by the sensor. A widely used iterative method to refine blurred images into sharper representations is the Richardson–Lucy deconvolution.^12–14^ This approximation iteratively recovers a sharper estimate of the object by comparing the observed blurred image with the modeled or measured point spread function (PSF) of the imaging system. While this method can significantly enhance resolution, it is computationally intensive, demands precise knowledge of the depth-dependent PSF, and often requires many iterations to converge to a high-fidelity reconstruction.

Artificial Intelligence (AI), particularly deep learning, has rapidly transformed the landscape of biological imaging.^15,16^ By learning patterns from vast datasets, AI models can infer or reconstruct high-quality images from limited or noisy input,^6^ offering a powerful alternative to traditional physics-based methods. In microscopy, AI has already enabled super-resolution reconstruction,^17^ image denoising,^6^ segmentation,^18^ and rapid volumetric imaging,^19^ often surpassing the speed and accuracy of classical approaches.^5^ These advances are especially impactful for techniques like LFM, where image formation is computationally demanding due to the complex mapping of angular and spatial information.^20^ By leveraging AI, such systems can now decode and reconstruct 3D biological scenes in real time, even under ultra-low-light conditions, as is the case for bioluminescent signals.

In this work, we present a Fourier light field microscopy (FLFM) framework combined with a deep learning–based reconstruction pipeline for rapid volumetric imaging under photon-limited conditions. Rather than aiming to recover a diffraction-unlimited representation of the sample, our approach is designed to invert the FLFM forward operator under realistic imaging conditions, including optical aberrations and low photon counts. To this end, we employ experimentally acquired wide-field (WF) z-stacks as reference volumes for supervised training, acknowledging that these do not constitute absolute ground truth due to the presence of out-of-focus contributions. Reconstruction quality is therefore evaluated in terms of structural fidelity relative to the reference modality and functional interpretability in biological applications. This strategy enables robust, real-time volumetric reconstruction in both fluorescent and bioluminescent regimes while maintaining consistency with the underlying imaging physics.

## Results and discussion

### Design and performance of the Fourier light field microscope

We first established the optical design parameters to achieve a depth of field of approximately 100 µm over a 150 µm field of view while maximizing the attainable spatial resolution. We built the instrument around a 40×/1.25-NA silicon oil-immersion objective to maximize photon collection with a high numerical aperture avoiding spatial oversampling with an illumination path for fluorescence excitation and transmitted brightfield imaging. The optical parameters of the microscope were carefully selected to meet performance criteria as calculated by classical reconstruction using light-field deconvolution. We used a microlens array (MLA) with a pitch of 1 *mm* and a focal length of 30.6 *mm* to target 2.1 *µ*m lateral resolution (*R_xy_*), 16.5 *µm* (*R_z_*), 88 *µm* (DoF), and 147 *µm* (FOV). We positioned the MLA in the conjugate back focal plane, and projected its image onto a single-photon-resolving qCMOS camera using a pair of relay lenses with f=100mm. Additionally, a 5-mm-diameter iris positioned in a plane conjugate to the native image plane of the objective acts as a field stop, restricting the spatial field of view and thereby preventing overlap between adjacent angular channels on the image sensor (Fig. 1a;^10^).

**Figure 1:**
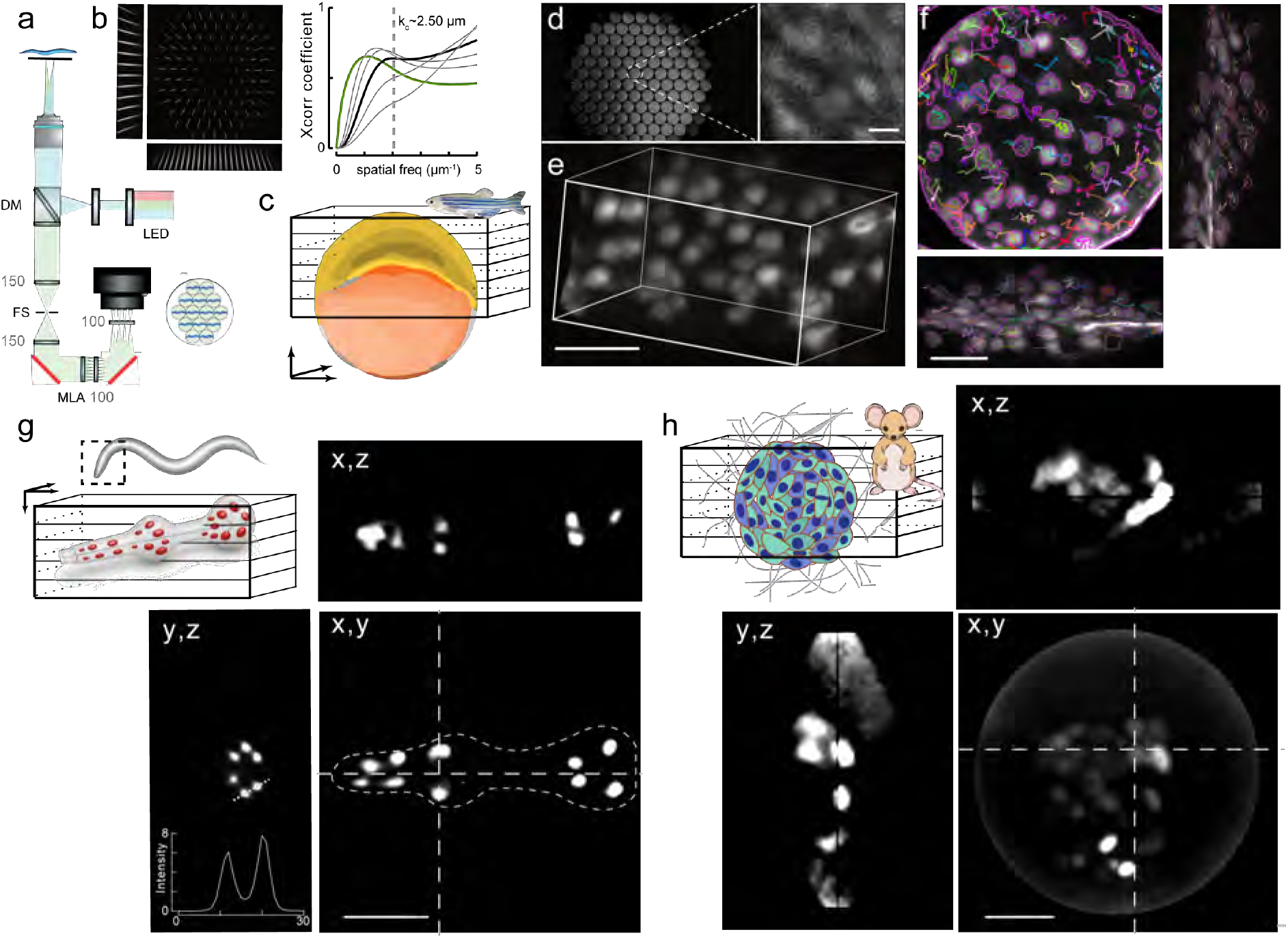
Single-exposure Fourier light field microscopy. **a.** Schematic of the Fourier light field microscope (FLFM). Grey numbers next to the elements indicate the focal length (in mm) of the depicted lenses; MLA, microlens array; FS, field stop; LED, light emitting diode. **b.** Spatial frequencies recovered from the PSF acquired with 200nm microspheres, using a parameter-free image resolution estimation method based in on decorrelation analysis. The upper inset shows the PSF of the system, , plot on the right shows some decorrelation functions computed for the image and resolution estimation. Black curve indicates decorrelation function of highest frequency peak, with a cutoff at 2.5µm. **c.** Schematic of a zebrafish embryo at gastrulation stage. The square indicates the imaged region. **d.** Raw FLF image showing 91 perspectives of the zebrafish embryo through the microlens array, with the central elemental image magnified. Scale bar = 10µm. **e.** 3D reconstruction at time 0 from sequence shown in Supplementary Video S1 using classical Richardson-Lucy deconvolution. Scale bar = 20µm. **f.** 3D segmentation of cells from a single plane of Supplementary Video S1, overlaid with their tracked trajectories across the X, Y, and Z projections. Scale bar = 20 µm. **g.** Representative examples of pharyngeal nuclei of transgenic *C. elegans*. Orthogonal views show the x,z and y,z section along the indicated lines. The intensity profile in the y,z view shows the signal distribution along the dotted line. **h.** Mouse stem cell colonies (5 days post-sampling) expressing a fluorescent protein in the nucleus, deconvolved with RL algorithm. Images display the maximum intensity projection along the z-axis, with side views shown along the indicated dotted lines. Scale bar = 50µm.

To evaluate the performance of the FLF microscope we assessed the PSF by imaging *d*=200 nm fluorescent microspheres and measured the highest frequency distribution in the 2D power spectral density. In agreement with the design parameters and the target size, the largest recovered frequencies correspond to the smallest size in the image (Fig. 1b). Across the central field, we estimated spatial frequencies (*k_c_*≈ 0.4µm*^−^*^1^, 2.5µm) according to^21^ consistent with our 2.1 µm target (Fig. 1b). As expected from the FLFM optical design, elemental images (EIs) acquired through different lens lets exhibited distinct but complementary information content. Through pairwise structural similarity analysis, we confirmed that this angular diversity is preserved in our experimental implementation (Supplementary Fig. S1). We next tested the imaging procedure across different fluorescently labeled biological specimens, including gastrulating zebrafish embryos imaged in short time-lapse sequences (30 min, 3-min intervals; Fig. 1c–f), *C. elegans* (Fig. 1g), and spheroids derived from mouse embryonic stem cells (mESCs; Fig. 1h). In all three systems, cell nuclei were fluorescently labeled using histone-GFP in zebrafish, Histone 2B-mTurquoise2-enhanced Nano-lantern (H2B-TeNL) in mESCs, and a GFP fused to a nuclear localization signal (NLS-GFP) in the pharynx of *C. elegans*, respectively. To assess time-resolved volumetric imaging, we focused on gastrulating zebrafish embryos and acquired light-field images every 3 min over a period of 27 min (Supplementary Video S1). To restore the volumetric from the plenoptic information we resorted to conventional Richardson-Lucy deconvolution algorithms.^10^ Although we were able to obtain high-quality 3D representation from plenoptic images (Fig. 1e), which facilitated single nuclei tracking in 3D over a volume of 140×140×100 µm (Fig. 1f), this classical deconvolution was computationally expensive and thus proved prohibitive for imaging longitudinal biological dynamics. After reconstructing the nuclear signal distribution in the *C. elegans* pharyngeal muscles, we resolved all 20 expected nuclei (Fig. 1g). Despite this overall faithful recovery of the nuclear organization, image deconvolution took five min per image and the orthogonal views revealed reconstruction artifacts, particularly along the axial dimension (Fig. 1g). Similar artifacts were apparent in the reconstruction of mESC spheroids (Fig. 1h). In optically heterogeneous or deeper specimens, sample-induced aberrations and scattering can cause the effective PSF to deviate from that assumed for Richardson–Lucy deconvolution,^22^ providing one possible source of such reconstruction artifacts. To address this, we sought to accelerate reconstruction with AI-based models.

### Attention-based Deep Learning for Light Field Reconstruction

Several deep learning methods have been introduced recently such as VCDNet,^20^ F-VCD,^11^ HyFLM and VsLFM^23^ and V2V3D,^24^ all focused on increasing resolution during fluorescent imaging with a high photon budget. VCD-Net established a learned reconstruction framework for volumetric fluorescence imaging from light-field data. HyLFM-Net instead used a hybrid light-field/light-sheet platform, where concomitantly acquired high-resolution light-sheet planes continuously served as training data and continuous validation/refinement targets during long time-lapse experiments.^25^ This was recently extended to incorporate channel attention (HyLFM-A-Net), to replace iterative tomography for fast reconstruction at the expense of robustness. VsLFM is a physics-based deep-learning framework trained on a large scanning-derived dataset to push LFM toward diffraction-limited performance. To increase generalization capability across diverse structures, VsLFM (virtual-scanning light-field microscopy) was introduced, a physics-based deep learning framework designed to improve light-field microscopy toward diffraction-limited resolution by learning the frequency-aliasing relationship between angular views from a large paired dataset built from scanning LFM measurements. V2V3D introduced an unsupervised view-to-view reconstruction strategy for light-field microscopy, splitting the angular views into two subsets and leveraging their cross-consistency to jointly denoise and reconstruct 3D volumes, with wave-optics-based feature alignment to better recover high-frequency detail.^24^

F-VCD is a two-stage VCD reconstruction strategy for fluorescence imaging with a Fourier Light field microscope.^11^ F-VCD decomposes Fourier light-field reconstruction into two sequential stages: a view-correlated denoising module that suppresses noise across the light-field views, followed by a limited-view 3D reconstruction module that maps the cleaned light-field data to a high-resolution volumetric output. None of the previously reported networks combine end-to-end denoising and reconstruction with the robustness of supervised learning under ultra-low-light conditions. Supervised deep learning is more reliable than self-supervision when signal levels approach the noise floor, but it requires paired experimental and reference images,^26^ which in our case were obtained by simultaneous wide-field stack and plenoptic light-field imaging.

To enable multimodal imaging, we incorporate WF emission detection along the FLF path (Fig. 2a). Fluorescence excitation with WF detection provided paired reference data as volumetric targets: a 3 dimensional WF stack and the corresponding plenoptic LF image of the same sample (Fig. 2a). This configuration enabled us to generate the paired WF and LF datasets required for supervised training.

**Figure 2:**
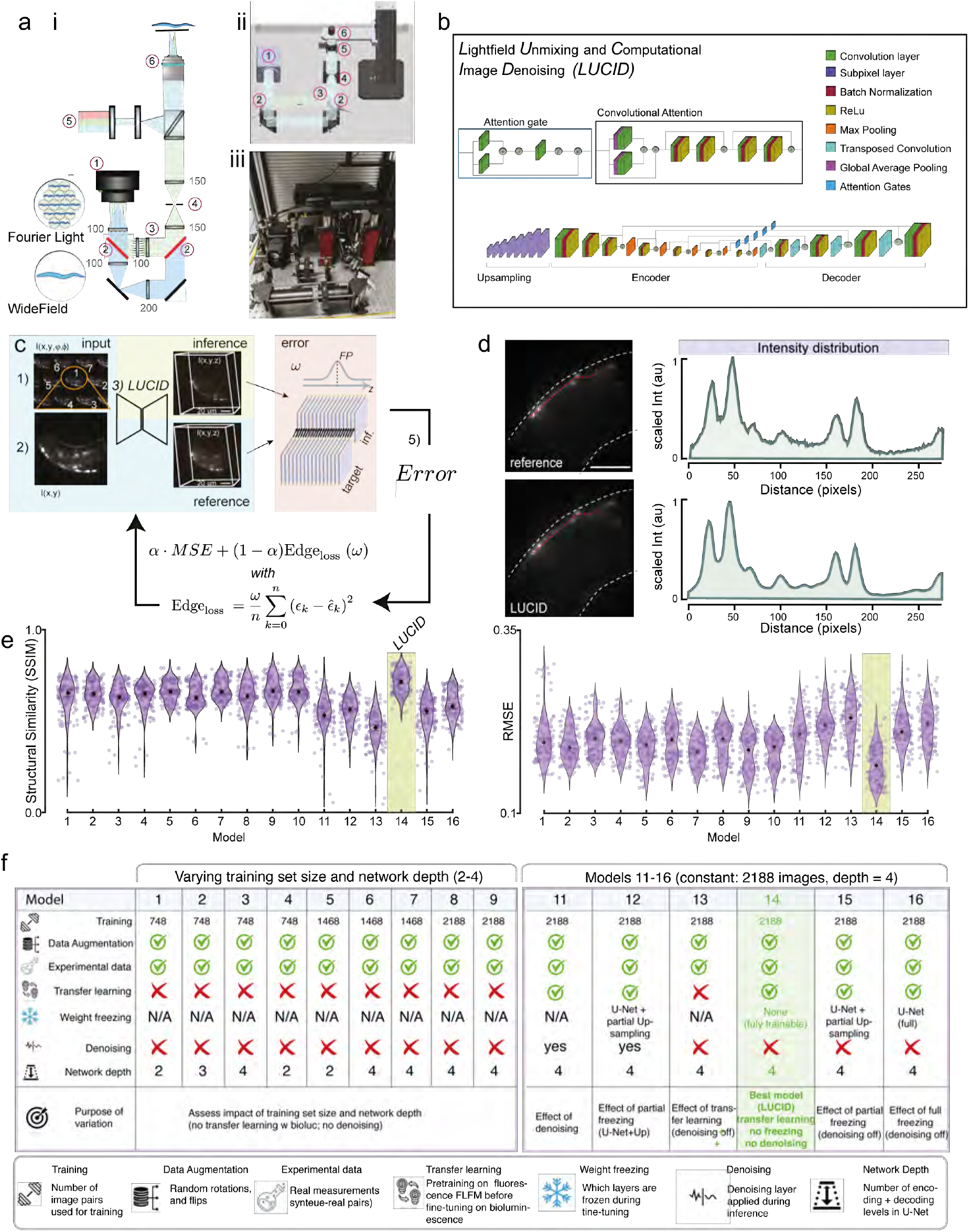
Deep learning pipelines for Multimodal Fourier light field microscopy. **a.** i) Schematic of the dual optical-branch system for sequential acquisition of WF and FLF images. The setup uses motorized flipper mirrors to dynamically switch the optical path between two branches: one configured for FLF imaging and the other for conventional WF imaging. Grey numbers next to the elements indicate the focal length (in mm) of the depicted lenses; 1) Camera, 2) motorized flip-mirror, 3) MLA, 4) iris, 5) LED, 6) objective lens. ii) 3D sketch of the setup. Numbers indicated as in (i). iii) Actual photograph of the setup. **b.** Schematic of LUCID (Light-field unmixing and computational image denoising), a pipeline for real-time volumetric reconstruction and signal restoration from noisy lightfield input data using an attention-based convolutional neural network (CNN) trained on synthetic and fluorescent imaging data. **c.** Training pipeline for FLF models in 3D reconstruction. The process consists of five steps: (1) data acquisition, (2) extraction of the elemental images, (3) feed-forward pass through LUCID, (4) comparison of predictions with the reference images, and (5) gradient calculation to refine performance in the next cycle. A custom loss function combining Mean Squared Error (MSE) with a Sobel-based edge loss was used to improve edge fidelity and reduce over-smoothing. The Sobel loss was applied across all slices of the 3D stacks with a Gaussian weighting scheme, *ω*, that emphasized the central, information-rich focal plane (FP, see Text and Methods for details). **d.** Intensity profiles measured along a line intersecting neuronal edges in *C. elegans* (red dashed line) on reference volume and the neural network reconstruction. Input image SNR = 3.648. Reconstruction metrics mSSIM = 0.73, mNRMSE = 0.129. Scale bar 20 *µ*m. **e.** Model performance was evaluated on unseen fluorescence data using SSIM and RMSE relative to the reference (N = 34 images/ model; N = 136 after augmentation). Sixteen network configurations were compared, with Model #14 (LUCID) achieving the highest similarity and lowest error and outperforming the second- (#) and third-best (*) models (p = 0.0032 and p = 0.001, respectively; Dunn’s post hoc test with FDR adjustment). **f.** Systematic evaluation of LUCID training parameters (Supplementary Table 1). Models 1–9 assess training-set size and network depth, whereas Models 11–16 evaluate denoising and transfer-learning strategies using a fixed dataset (2,188 image pairs) and network depth of four. Model 14, combining denoising and transfer learning with full network fine-tuning, showed the best performance and was selected as the final LUCID model. Symbols indicate the parameters varied across models.

Artificial neural networks have proven to be effective in accelerating volumetric reconstructions of LF microscopy, overcoming the computational bottlenecks of traditional deconvolution approaches.^5^ However, standard convolutional neural networks (CNNs) apply the same convolutional kernels across the image, which may limit their ability to adapt to context-dependent local structures, which can lead to insufficient adaptation to local structures, potentially amplifying noise or blurring fine details in the reconstruction.^27^ To address this, we developed and deployed a attention-based convolutional neural network optimized for Fourier light field (FLF) reconstruction.

Our base model is built on a U-Net backbone and integrates two attention components: attention gates and convolutional block attention (Fig. 2b, ref.^28^). Attention gates refine skip connections by selecting only the most relevant spatial information, while convolutional attention blocks, placed within the encoder, preserve high-frequency features typically lost during downsampling.^28^ By integrating attention mechanisms, the network can dynamically weigh the importance of different spatial features, allowing it to focus on informative regions while suppressing irrelevant noise and artifacts.^29^ This selective enhancement improves the fidelity of the reconstructed images by preserving fine structural details and reducing background interference.

Previously, we have shown that denoising input data before feeding them into the network improves reconstruction metrics,^5^ which required two separate networks: a content-aware restoration network^6^ and a VCD network.^20^ This approach not only increased the computational cost, but also substantially increased workload and time, as it required two independently trained networks-one for denoising and one for reconstruction.^5^ Likewise, *rad* - FLFM sequentially combines different algorithms for denoising and reconstruction,^30^ which propagates errors across processing steps, increases computational complexity, and prevents the end-to-end optimization of image restoration and volumetric reconstruction. Recent approaches such as V2V3D leverage view-to-view consistency to jointly perform denoising and volumetric reconstruction in a self-supervised manner. These methods rely on the assumption that signal is consistent across angular views, while noise is uncorrelated. While effective in moderate-signal-to-noise ratio (SNR) fluorescence regimes, such assumptions may break down in photon-starved bioluminescence imaging, where signal levels approach the noise floor and inter-view consistency is reduced. In contrast, our end-to-end network is trained directly on both noisy input projections and high-SNR reference volumes, enabling it to simultaneously learn image reconstruction and noise suppression. The experimental training dataset was carefully designed to include both low-SNR and high-SNR images of the same ROI, achieved by intentionally reducing the laser power and exposure time for the low-SNR conditions. This behavior emerges from the structure of the data and the architecture design, rather than from explicit constraints or losses. The complementary angular information encoded by each FLFM lenslet (Supplementary Fig 1) allows every elemental image to be treated as an independent training sample, providing a scalable source of experimental data augmentation that enhances denoising performance while minimizing redundancy and overfitting. As a result, the network performs reconstruction and noise attenuation simultaneously, without introducing additional complexity to the pipeline. Consequently, our model improves reconstruction stability relative to models without implicit denoising (Supplementary Fig. S2), eliminates the need for preprocessing or auxiliary networks, and enables robust volumetric inference under ultra-low-SNR conditions.

Furthermore, we used a custom loss function that combines root mean squared error (RMSE) with a Sobel-based edge loss^31^ to regularize gradient consistency and mitigate oversmoothing leading to edge fidelity in the predicted output (Fig. 2c). While the MSE captures pixel-wise differences between the predicted and the reference stacks, the Sobel component emphasizes discrepancies in edge information, helping the network better learn spatial structures. To further refine this, we applied the Sobel loss across all slices of the 3D stacks but introduced a Gaussian weighting scheme that prioritizes the central slices, the focal plane, where most of the relevant information is typically concentrated (Fig. 2c). This weighting ensures that errors in the focal region are penalized more heavily than those in peripheral slices, guiding the network to learn from the most informative regions of the data and resulting in reconstructions that preserve the intensity distribution of the experimental reference modality without introducing detectable nonlinear distortions (Fig. 2d).

To train the network, we generated synthetic images by forward projection of highresolution wide-field stacks using wave optics-based PSFs,^5,20^ yielding *N* = 2188 pairs of images. While synthetic pre-training aided convergence, artifacts emerged from mismatches with real optics, leading to degraded performance (Fig. 2e, f). To enhance the network’s relevance to real-world applications, we combined synthetic and experimental FLF data, in a three-stage strategy: pretraining with synthetic data, fine-tuning with experimental fluorescence images derived from *C. elegans* labeled with fluorescent protein in their muscles and neurons, and final training with bioluminescent data with the same expression pattern. We developed custom software to automatically extract 91 angular perspectives (200 × 200 pixels each) from raw FLF images (Fig. 2c). The pipeline supports varying microlens array configurations and aligns each input with its corresponding wide-field stack, acquired from the same sample using a motorized flip mirror system (Fig. 2a). The reference stacks were augmented through flipping, rotation, and *z* inversion to improve robustness. Each WF volume spans 101 axial planes over a 100 µm depth and in total, we obtained *N* = 2000 paired datasets from our experiments. For each combination, we trained and evaluated different models by measuring structural similarity (SSIM)to quantify consistency with experimentally acquired reference data, RMSE (Fig. 2f, Supplementary Table 1), and resolution by means of the power spectral density across reconstructions of unseen samples. In total, we compared 16 models (Fig. 2e) and we named the best-performing model *LUCID* (*L*ightfield *U* nmixing and *C* omputational *I* mage *D* enoising). LUCID consistently suppressed noise, generalized across datasets (see validation section below), and preserved structural details and spatial resolution and achieved up to ≈ 90× faster reconstruction than our GPU-accelerated RL baseline on matched datasets (Supplementary Fig. S3), without introducing artefacts such as step-like structures or blodges when reconstructing an intensity gradient or purely noisy input data (Supplementary Fig. S4).

### Cross-validation and domain adaptability

To demonstrate the adaptability and transferability of the LUCID base model, we finetuned the network for diverse biological specimens and imaging modalities and validated its performance on fluorescent body wall muscles (BWM) and nuclear morphodynamics in *C. elegans*, as well as mouse embryonic stem cells, bioluminescent nuclear imaging in mESCs, and bioluminescent calcium imaging in muscles and neurons.

### Imaging of nuclear dynamics in moving animals

In conventional 3D time-lapse microscopy, such as confocal or light-sheet microscopy, imaging an extended depth of field requires sequential acquisition of multiple focal planes at rates sufficiently fast to avoid motion artifacts during volumetric imaging. Because *C. elegans* is a highly motile organism, capturing subcellular dynamics in three dimensions remains a substantial technical challenge.

Our FLFM approach addresses this limitation by enabling sub-second volumetric imaging in a single exposure, thereby eliminating the inter-slice inconsistencies inherent to z-stack acquisition in moving specimens while maintaining the native lateral resolution of the FLFM system. To demonstrate this capability, we sought to resolve tissue deformations in freely moving animals using GFP::LMN-1-labeled nuclei as fiducial markers (Fig. 3a–c; Supplementary Videos S2 and S3).

**Figure 3:**
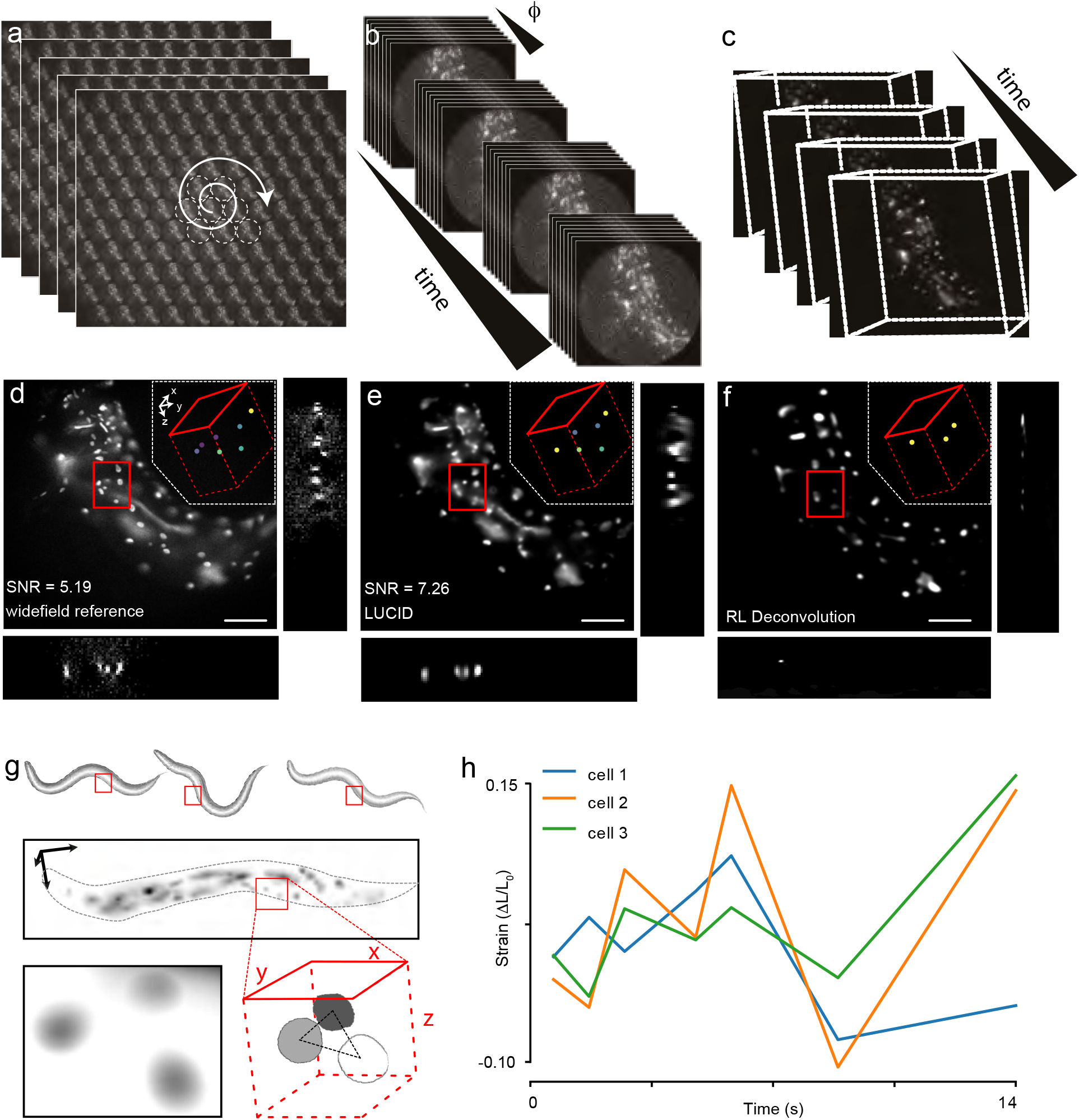
Nuclear dynamics in freely moving worms. **a–c.** Schematic of the image-processing workflow. (a,) Time series of raw Fourier light-field images acquired from a freely moving animal. (b,) Angular views are extracted from each light-field image and processed by LUCID to generate a time series of volumetric reconstructions. (c,) Recon-structed 3D volumes enable tracking of fluorescent structures over time.**d–f,** Representative volumetric reconstructions obtained from the wide-field reference stack (d), LUCID (e), and Richardson–Lucy (RL) deconvolution of the corresponding light-field data (f). Orthogonal projections of the region indicated by the red box illustrate axial localization of individual fluorescent structures; insets show their spatial distribution within the reconstructed volume. Note the missing structures in the the inset of (f). SNR values are indicated for the wide-field reference and LUCID reconstruction. **g,** Application of LUCID to track fluorescently labeled nuclei during freely moving *C. elegans* locomotion. Schematics indicated the approximate location of identified nuclei. Individual cells are localized in three dimensions and their relative displacement is used to quantify local tissue deformation. **h,** Representative time courses of strain, (Δ*L/L*_0_), determined from the relative positions of three tracked cells over 14 s. See also Supplementary Fig. S5 for the stationary-animal control used to estimate the strain noise floor.

This application also provided an opportunity to evaluate the adaptability of LUCID to a new imaging regime. Specifically, LMN-1::GFP-labeled nuclei were imaged using a 20x objective rather than the 40x objective used for the initial training, introducing substantial changes in sample morphology, spatial scale, and the angular encoding of the Fourier light field data. As expected, the original model exhibited reduced reconstruction performance under these conditions. Fine-tuning the network on a modest dataset (Supplementary Table S2) acquired with the new imaging configuration restored reconstruction fidelity, demonstrating that LUCID readily adapts to changes in specimen type and optical configuration without modification of the underlying network architecture.

Under photon-limited imaging conditions, LUCID produced smoother and more temporally stable volumetric reconstructions than RL deconvolution. Because experimentally accessible volumetric ground truth is unavailable under these imaging conditions, reconstruction fidelity cannot be rigorously validated against an optically sectioned reference. Nevertheless, LUCID reconstructions showed higher agreement with the WF reference modality based qualitative comparison of cell localization (Fig. 3d-f). We then used short timelapse videos to segment and track individual nuclei in 3D of freely moving animals (Fig. 3g; Supplementary Video S3). To obtain the local tissue deformation, we obtained the nearest-neighbor distance between selected nuclei (Fig. 3h), and calculated the change over time. This strain map indicates that cells are subject to both compression and extension, which brings nuclei closer or farther away from each other. An animal that was anesthetized and therefore did not move during the acquisition showed that the calculated strain is capturing tissue movements and does not reflect tracking/reconstruction uncertainties (Supplementary Fig. S5). Together, these results demonstrate that light field imaging combined with LUCID restoration enables 3D segmentation and quantitative strain mapping of subcellular dynamics in freely moving animals, while minimizing motion artifacts relative to z-scanning.

### Cellular dynamics within mESC spheroids

To assess whether the LUCID base model can be efficiently adapted across species and imaging contexts, we apply our LUCID model originally trained on fluorescent *C. elegans* nuclei and to spheroids derived from mouse embryonic stem cells expressing fluorescent nuclear markers. Despite the increased cellular density and distinct tissue organization, the retrained model accurately resolved individual nuclei throughout the spheroid volume (Fig. 4a–c; Supplementary Video S4). Reconstruction of the complete 60-minute time-lapse sequence (120 volumes, 0.5 MP per frame) was completed in approximately 1 minute, compared to 1.5 hours using Richardson–Lucy deconvolution on the same workstation (Supplementary Fig. S3). Furthermore, LUCID substantially increased SNR compared to the reference image and similar to RL deconvolution (Fig. 4d), facilitating nuclear segmentation and tracking. In the absence of a parallel accessible volumetric ground truth (due to intrinsic sample motility), performance was therefore assessed based on the preservation of biologically meaningful downstream measurements. Therefore, we used these volumes and we quantified nuclear trajectories, nearest-neighbor distances, and mean squared displacements in three dimensions, revealing heterogeneous and superdiffusive nuclear dynamics within the developing spheroid (Fig. 4e–h).

**Figure 4:**
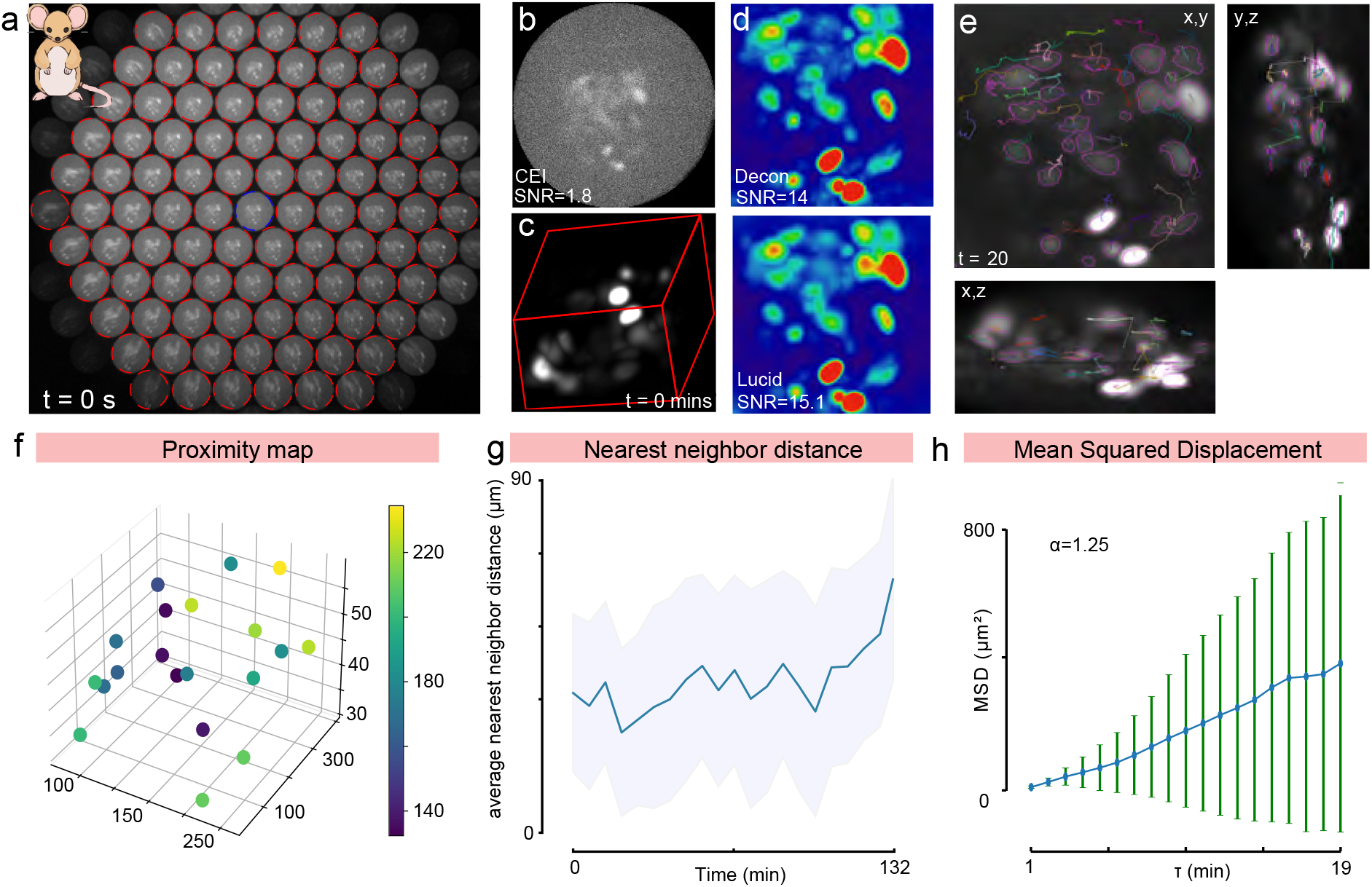
LUCID enables 3D segmentation and tracking of mESC spheroids with nuclear fluorescence. **a.** Raw FLF image, with all automatically detected perspectives in red outlines, and the central in blue. **b.** The central elemental image (CEI, highlighted in blue in a) showing the raw image of the mESC spheroid and the associated signal/noise ratio (SNR). **c.** 3D volume of the reconstructed light field image at the initial time point (t = 0 min). See also Supplementary Video S2. **d.** Representative comparison of volumetric reconstructions obtained by Richardson–Lucy deconvolution and LUCID. SNR=signal to noise ratio. **e.** Orthogonal projections (Z, X and Y) of the reconstructed spheroid after 20 min. Colored trajectories indicate the three-dimensional displacement of individual cells over time. **f** Three-dimensional proximity map of tracked cells, color-coded according to nearest-neighbor distance. **g.** Average nearest-neighbor distance of tracked cells over 132 min; shaded region indicates the variability (mean±SD) across cells. **h.** Mean-squared displacement (MSD) of tracked cells as a function of lag time (*τ*). The scaling exponent (*α*=1.25) derived from a fit to a power law indicates superdiffusive cellular motion. Error bars indicate variability across tracked cells.

After 3D segmentation, we calculated the proximity map, which indicates the nearest cell neighbors (Fig. 4f) for every frame in the reconstructed image stack. We found that the average nearest-neighbor distance increases, concomitant with the growth of the cell cluster (Fig. 4g). This reveals how cells are packed in tissues and facilitates to study growth, (un)jamming transitions, cell sorting and migration, or tissue tension.

Next, we tracked each segmented nucleus and calculated the mean squared displacement (Fig. 4h). Power-law fitting of the MSD vs time lag yielded a scaling exponent *α*=1.25,

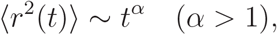

consistent with super-diffusive, directed motion rather than a random walk. This indicates that cells actively migrate apart, leading to reduced packing density and a transition of the spheroid from a more compact, solid-like state toward an expanding tissue during growth. Together, this shows that the single exposure 3D imaging reveal changes in tissue architecture in a nearly noninvasive fashion.

### Excitation-free imaging of biological dynamics with bioluminescence

Because *C. elegans* animals have photoreceptors and avoid even low intensities of blue and orange light,^32,33^ conventional fluorescence imaging causes frequent reversal, escape responses that obscure the natural behavioral repertoire.^34^ Likewise, calcium imaging using GCaMP family sensors require blue light excitation, which activates neurons in photosensitive animals such as *C. elegans*. This response is often stronger than the response to other physiological stimuli.^35,36^ Bioluminescence imaging is an attractive alternative because it does not require external excitation light. However, due to the low photon yield of many luciferases, conventional bioluminescence microscopy relies on long exposure times, leaving fast volumetric imaging of subcellular biological dynamics out of reach so far. We recently showed that light-field microscopy marries well with photon-starved bioluminescent samples, due to their ability to capture volumetric information in one single exposure.^5^ This approach reduces acquisition time by eliminating the need to scan through multiple planes, as required in conventional confocal or WF z-stacks, thereby preserving the frame rates necessary to capture rapid biological events such as calcium dynamics and stress responses. The tradeoff, however, is reduced spatial resolution, which has so far limited our ability to record physiological signals from individual neurons.^5^ Here, we extend this approach to reconstruct 3D image stacks in real time using bioluminescence Fourier light field microscopy. The training pipeline. The LUCID models initially trained on fluorescence data recovered the overall sample morphology but struggled to reconstruct fine details in bioluminescent images due to the substantially lower photon budget and reduced SNR. To overcome this limitation, we generated a dedicated bioluminescence training dataset comprising paired low- and high-SNR FLFM images acquired at different exposure times (typically 1 s and 5 s), together with corresponding bioluminescent widefield z-stacks as reference volume (Figure 5a,b).

**Figure 5:**
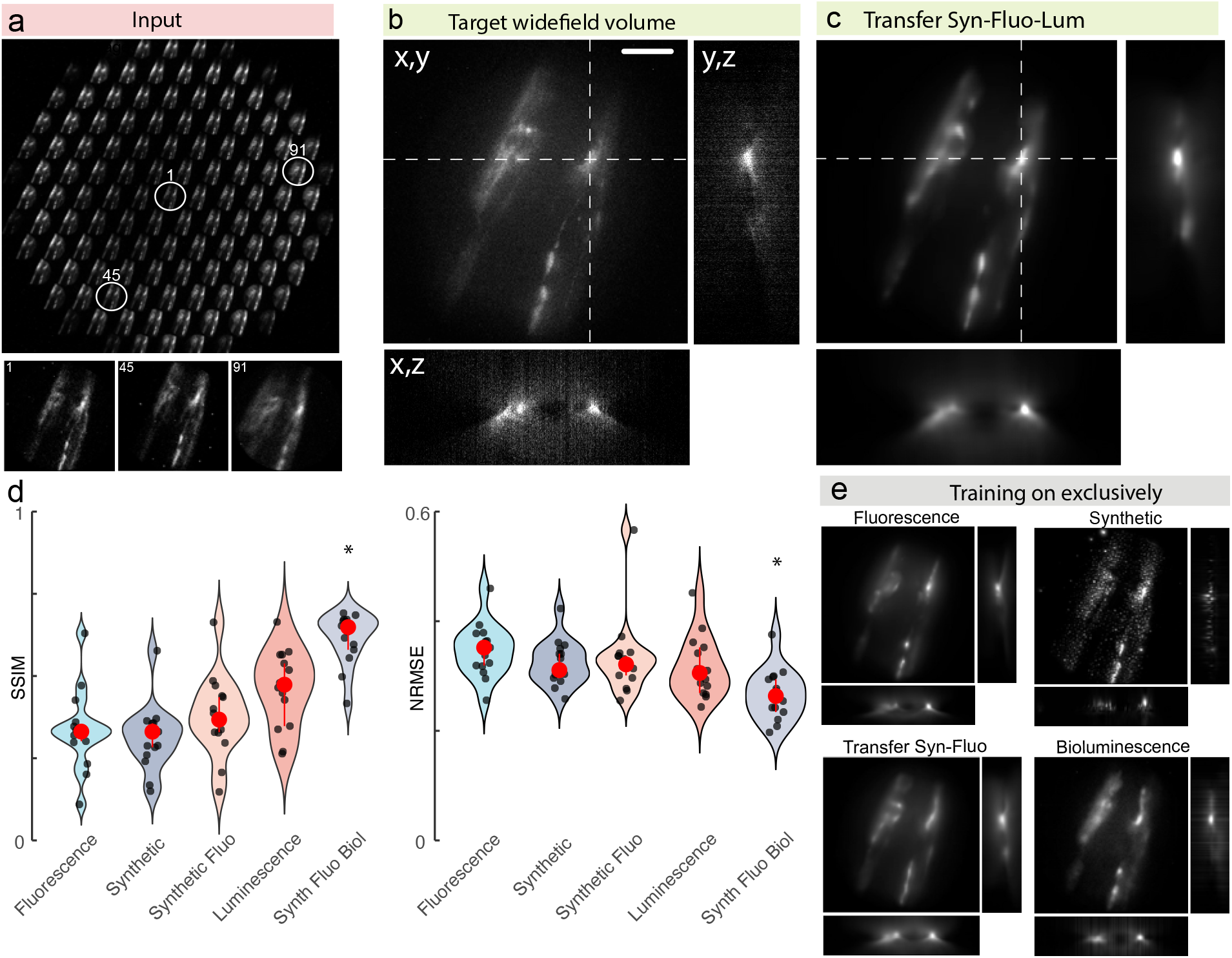
Transfer learning enables volumetric reconstruction of bioluminescence light-field data. **a.** Representative Fourier light-field input image showing the angular views of a bioluminescent labelled *C. elegans*. Selected views (1, 45 and 91) are shown below. **b.** Corresponding experimentally acquired wide-field reference volume shown as orthogonal (xy), (xz), and (yz) projections. **c.** Representative reconstruction obtained after sequential transfer learning from synthetic to fluorescence and bioluminescence data. **d.** Performance evaluation of individual models on unseen data using SSIM and NRMSE metrics (N = 15 images per model, no augmentation). Black dot, mean± standard deviation. The best-performing model is marked with an asterisk (*), showing a statistically significant improvement over the second-best model (p *<* 5.78e-06, two-sided t-test). Input image SNR = 3.029. Scale bar: 50 *µ*m.**e.** Representative volumetric reconstructions obtained using networks trained on exclusively fluorescence or synthetic data, and following sequential transfer learning from synthetic to fluorescence or to bioluminescence data. Orthogonal projections illustrate the corresponding axial reconstruction.

The best reconstruction performance was achieved using a three-stage transfer learning strategy consisting of pretraining on synthetic data, followed by fine-tuning on fluorescence and subsequently bioluminescence experimental datasets (Figure 5c, d). In contrast, models trained exclusively on synthetic, fluorescence, or bioluminescence data alone (Figure 5d, e) exhibited reduced performance, frequently failing to recover fine structural details or reproducing noise present in the training data. These results are expected due to different photon statistics and the absence of autofluorescence with bioluminescence contrast, highlight the importance of combining synthetic priors with progressive domain adaptation to achieve robust volumetric reconstruction of photon-limited bioluminescent samples.

Validation for functional bioluminescence imaging. To evaluate whether FLFM/LUCID reconstructions preserve spatially separable neuronal activity under photon-starved conditions, we performed experiments using animals expressing a bioluminescent calcium indicator under the control of a panneuronal (*rab-3* p) promotor. We observed calcium fluctuations in the motorneurons of the ventral nerve chord in freely moving animals (Supplementary Video S6, S7) and in the dorsorectal ganglion (Fig. 6a) of an immobilized animal. We recorded spontaneous activation of a neuron that we identified as DVA (Supplementary Video S8), and could visually separate neighboring neuronal structures approximately 5 µm apart (possibly DVC and DVB, Fig. 6a). After reconstruction, we observed more spatially confined signal distributions, and isotropic intensity profile in all three dimensions (Fig. 6b i-iii). We also observed that the spatial frequency content in the LUCID reonstructions is higher compared to the central elemental image, indicating an increase in resolution due to the training on high-resolution widefield data (Fig. 6c). After reconstruction, we extracted the calcium signal for the projected volume and compared it to the intensity extracted from the central elemental image (a coaxial ‘widefield’ view). We note that the central elemental view should be interpreted as an internal comparator extracted from the same acquisition, not as an absolute reference measurement. We then verified that reconstructed intensities preserve linear relationships with the reference signal (Supplementary Fig. S7), confirming suitability for functional imaging. In particular, temporal dynamics and relative amplitude variations in calcium signals were preserved after reconstruction, indicating that the model maintains quantitative relationships in the data. This is critical for downstream biological interpretation, where intensity linearity is required for accurate measurement of physiological activity.

**Figure 6:**
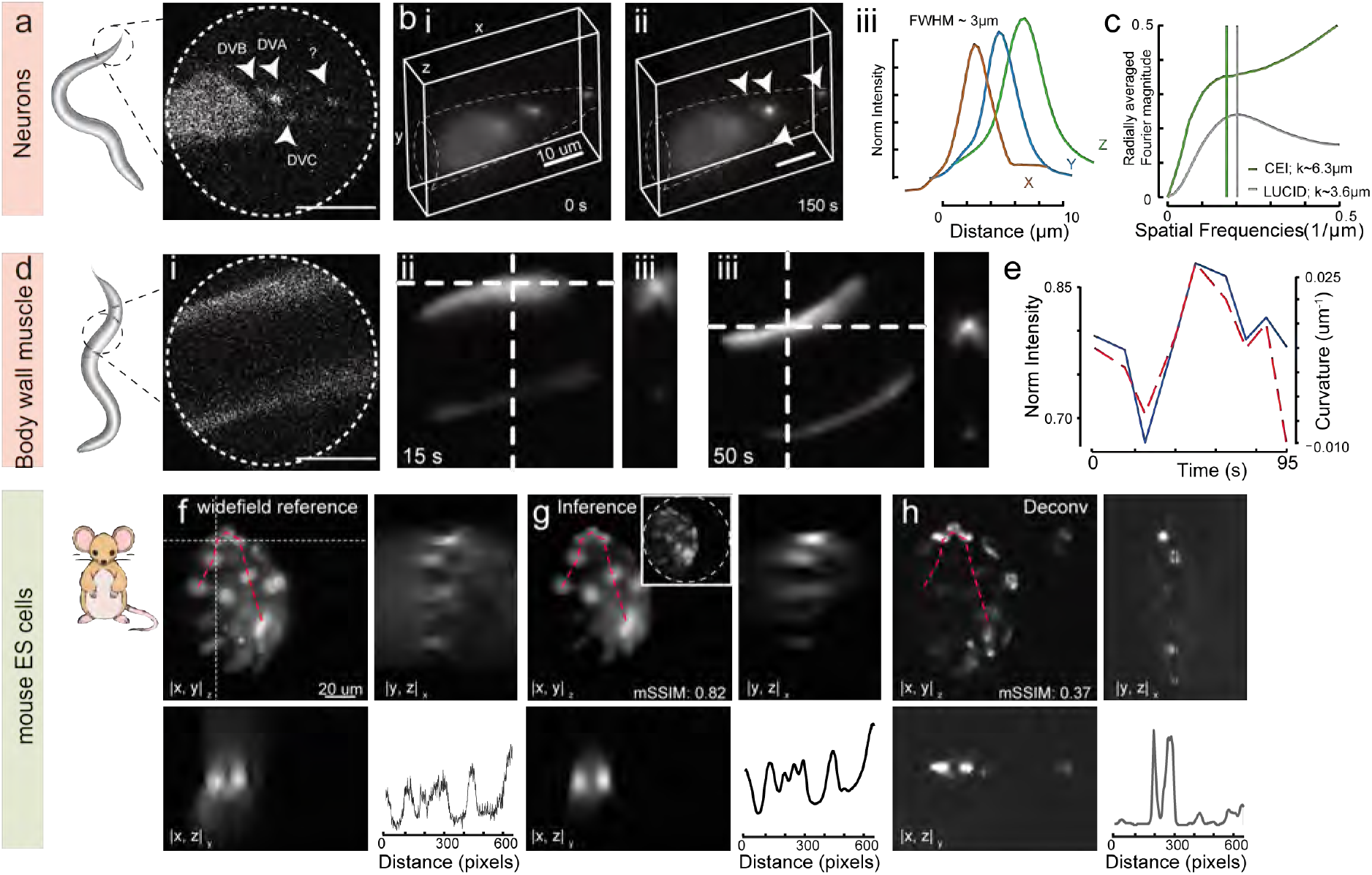
Validation of spatial and temporal information in LUCID reconstructions using bioluminescent input images. **a-c.** Volumetric reconstruction of luminescent neurons in *C. elegans*. (a) Representative Fourier light-field input image showing the DVA, DVB and DVC neurons; **b.** LUCID reconstructions at 0 and 150 s, respectively. Arrowheads indicate identified neuronal structures. (iii) Normalized intensity profiles through a representative reconstructed structure along the (x), (y), and (z) directions, showing a full width at half maximum (FWHM) of approximately 3 µm. **c.** Decorrelation function as a function of spatial frequency for the central elemental image of raw FLFM image and the corresponding LUCID reconstruction as an estimate for spatial resolution. **d.** Dynamic volumetric imaging of fluorescently labelled body-wall muscle (BWM) in *C. elegans*. i, representative Fourier light-field input; ii–iii, orthogonal views of LUCID reconstructions at 15 and 50 s, respectively. Dashed lines indicate the positions used to extract the corresponding orthogonal sections. **e.** Temporal evolution of normalized fluorescence intensity and body curvature extracted from the reconstructed volumes over 95 s. **f–h** Comparison of volumetric reconstructions of fluorescently labelled mouse embryonic stem cells. (f,) Experimentally acquired wide-field reference volume; (g,) LUCID inference from the corresponding Fourier light-field image; and (h,) Richardson–Lucy deconvolution of the Fourier light-field data. (xy), (xz), and (yz) projections are shown together with intensity profiles extracted along the indicated trajectories.

Next, to demonstrate the ability of our FLFM/LUCID combination to image photon-starved bioluminescent samples with short exposure times (≈1 s), we first imaged partially constrained *C. elegans* expressing a calcium-sensitive Nanolantern in body wall muscles (Fig. 6d). As stated above, we trained the network directly with bioluminescence data. This adaptation aimed to improve the network’s ability to handle low-quality inputs while still producing high-quality reconstructions. Consequently, the 3D reconstruction of the bioluminescence scene was achieved seamlessly through this single-network pipeline, eliminating the need for a separate denoising network

As previously described, we observed that the intensity of bioluminescence is correlated with curvature during body bending,^5^ indicative of higher calcium concentrations during muscle contractions (Fig. 6e).

Validation on bioluminescent stem cell spheroids. Finally, we used our LUCID pipelines to reconstruct light-field images recorded from bioluminescent mESC derived spheroids. In contrast to sparse neuronal samples, densely packed cell aggregates present additional challenges due to low per-cell photon counts, increased optical scattering, and reduced contrast in the raw data.

Existing models trained on fluorescent Histone 2B-labeled or bioluminescent neuronal datasets did not generalize reliably to this new domain, frequently reconstructing individual nuclei as clusters, likely due to differences in spatial scale and cellular organization (not shown). To account for these differences, we fine-tuned the pre-trained LUCID model using a dedicated bioluminescent stem-cell dataset. Because the network had already learned general features relevant to volumetric reconstruction, adaptation required only a modest amount of additional training data (10 GB). Fine-tuning was completed in approximately one hour on a single GPU and substantially improved the separation and localization of individual nuclei. Comparison of spatial intensity profiles between the WF reference, LUCID reconstruction, and Richardson–Lucy deconvolution further showed that LUCID closely preserved the position and relative intensity distribution of individual nuclei observed in the experimental reference volume (Fig. 6f–h). These results show that LUCID can be efficiently adapted to new biological specimens and imaging conditions through transfer learning without modifying the underlying network architecture.

### Interpretability of the Fourier Light Field Reconstruction Network

After applying the network to biological reconstructions, we analyzed how LUCID derives 3D volumes from raw FLF images using Grad-CAM (Fig. 7), guiding the interpretabilty of the machine learning process by visualizing model attention in both, fluorescence and bioluminescence applications.^37–39^

**Figure 7:**
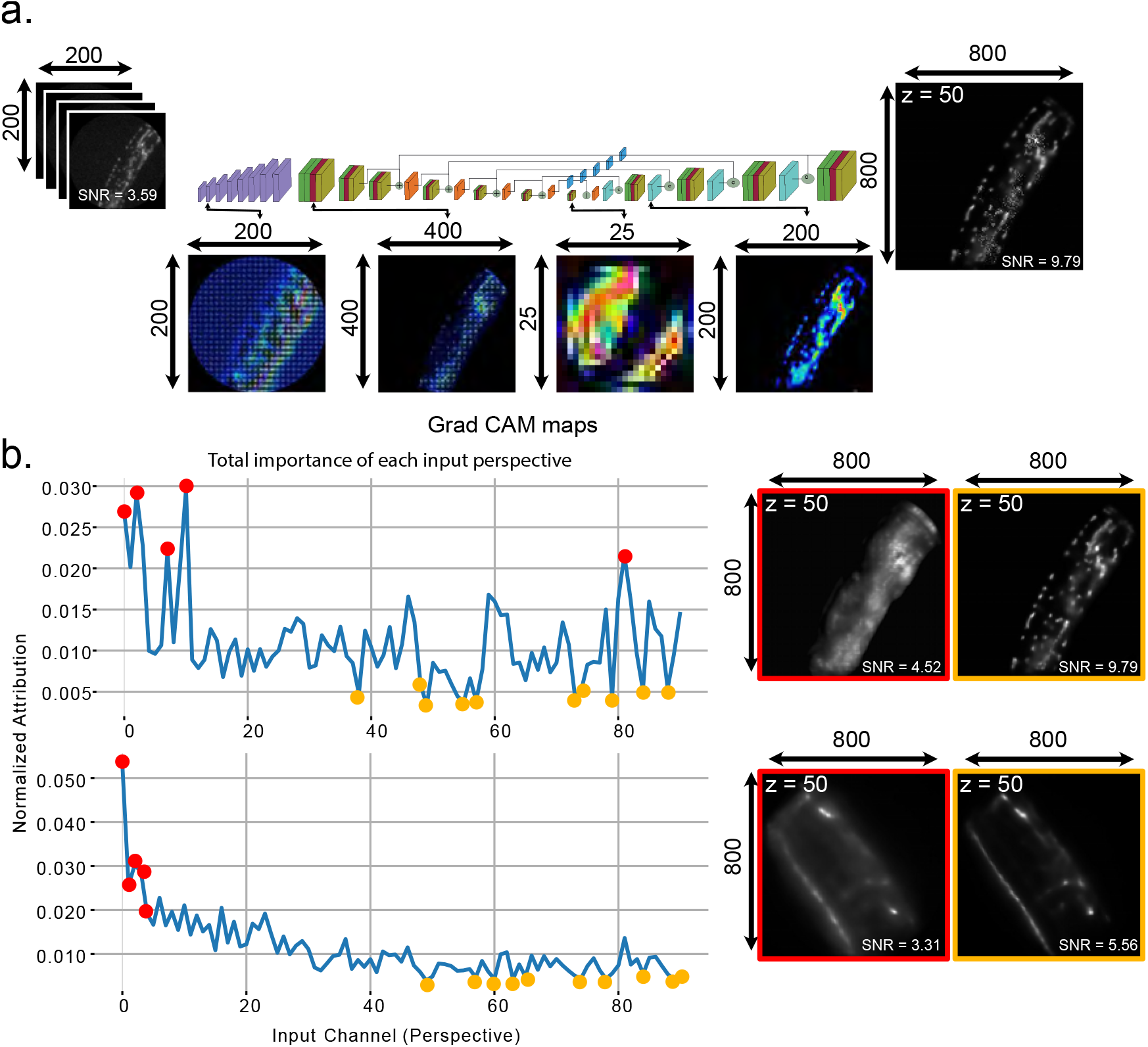
Interpretability analysis of Fourier light field reconstruction network. **a.** Grad-CAM spatial attention maps extracted at various stages of the U-Net, including input, bottleneck, and decoder layers, reveal how the network progressively learns to attend to salient spatial regions in the input. Heatmaps correspond to the reconstruction of the output slice at *z* = 50, the most information-rich plane. While input and decoder stage maps clearly highlight object structure, the bottleneck layer exhibits diffuse and spatially abstracted activations due to latent compression and feature generalization. Images depicted correspond to the *C. elegans* GFP::*lmn-1* strain. **b.** Channel-wise attribution analysis showing the relative contribution of each input perspective (elemental image) to the reconstruction, based on gradient magnitude. Two different networks trained on distinct datasets: fluorescence nuclear data (upper panels) and bioluminescence *rab-3* neuron data (lower panels). Each with unique attribution profiles. Red and yellow markers indicate the five most and ten least influential views, respectively. Reconstructions with red and yellow frames correspond to versions where these important/unimportant views were ablated. Removal of the most influential perspectives leads to severe loss of high-frequency content, while removal of the least important ones has negligible effect, validating the network’s learned prioritization of angular information.

### Investigating Spatial Attention Patterns in the Reconstruction Network

We examined attention maps in different stages of the U-Net: from the initial upsampling layers, through the encoder’s second convolutional block, the central bottleneck, and the first decoder block. These maps reveal how the model progressively shifts from diffuse low-level attention in the early stages to more focused, structured regions that align with key features in the output volume (Fig. 7a).

The central “bottleneck” layer behaved differently: its attention maps were more abstract and less localized. This is expected as this layer compresses the entire image into a compact internal representation that captures overall structure and context, rather than fine detail. Importantly, the network design includes skip connections that pass spatial detail from earlier layers directly to later ones, allowing it to combine both global context and local image features.

These findings suggest that the network relies on spatial regions associated with biologically relevant structures during reconstruction in a way that mirrors traditional image analysis - first parsing general features and then refining detail. Together, this interpretability analysis fosters confidence and provides practical insight for evaluating emerging reconstruction methods.

### Angular Attribution and View Selection

Having shown that all views in principle carry distinct information content (Supplementary Fig.1), we aimed to understand now how different angular views contribute to volume reconstruction, we computed gradient-based attribution scores for each input channel (elemental image). Central views — near the middle of the FLF layout — consistently showed higher influence, while peripheral views contributed less, likely due to lower signal quality or redundancy (Fig. 7b).

To validate this, we performed *in silico* ‘ablation experiments’. Removing the 5 most influential views led to a noticeable loss of fine detail, whereas removing 10 of the least important ones had minimal impact on reconstruction quality. This indicates that the network selectively leverages key perspectives for accurate 3D recovery and that attribution analysis can help optimize data acquisition, training pipelines and thus overall performance by identifying less influential views.

Together, these interpretability analyses provide a way forward to optimize Fourier light field imaging with reduced perspectives, which aids reconstruction fidelity without compromizing performance.

In summary, our results establish bioluminescence FLFM, catalyzed by our LUCID pipeline, as the a powerful approach capable of capturing fast, volumetric bioluminescence signals at near-single-neuron resolution, overcoming the long-standing trade-off between sensitivity and spatiotemporal resolution in photon-starved samples.

### Limitations

We noticed previously that the performance of the deep learning reconstructions is sensitive to input signal quality.^5^ We therefore tested the ability of LUCID to reconstruct input images with varying signal-to-noise ratio (SNR). We observed a clear performance drop below a specific value, which was quantified using SSIM against reference data. For fluorescence imaging, the effective lower limit is SNR ≈ 2.2 (Supplementary Fig. S6), while for bioluminescence imaging it is approximately SNR ≈ 0.3 (Supplementary Fig. S8). Inputs below these thresholds result in poor reconstructions as the model cannot reliably distinguish signal from noise.

In our optical design, we prioritized the depth of field to better capture the volume of the sample. Increasing the number of microlenses along a given axis extends the depth of field by sampling more angular information, but this comes at the expense of lateral resolution because of reduced spatial sampling. In future designs, this trade-off between depth and resolution could be mitigated by reducing the number of lenslets, thereby improving spatial detail while still maintaining sufficient depth coverage. The consequence of this limitation can be seen in LMN-1::GFP-expressing *C. elegans* (Fig. 3), where the nuclear envelope is poorly resolved because the intensity distribution is undersampled.

Our model also requires training on representative datasets to achieve optimal performance.While gradient-based regularization improves structural fidelity under low-SNR conditions, it may introduce bias toward sharper features in some contexts. Future work will explore alternative regularization strategies with explicit physical constraints. Our training pipeline involved conventional diffraction-limited epifluorescence or bioluminescence widefield stacks as reference volumes including residual out-of-focus signal, which may bias similarity-based evaluation metrics; thus, the reconstruction may be further improved using confocal or superresolved reference volume as ‘ground truth’ reference targets. Although transfer learning pipelines were implemented to ease this process, and we provide a ready-to-use Docker container, training remains a necessary step for domain adaptation to new biological samples or imaging conditions.

## Conclusion

In this work, we developed and validated a Fourier light field microscope (FLFM) integrated with a deep learning reconstruction pipeline, LUCID, to address practical limitations associated with volumetric imaging under photon-starved conditions of biological samples. Our optical design prioritized depth of field and field of view while maintaining micrometer-scale spatial detail, enabling single-exposure volumetric acquisition across diverse biological systems. Although classical deconvolution provided accurate reconstructions, it was prohibitively slow for longitudinal imaging. LUCID addressed this bottleneck by combining attention-based architectures, and custom loss functions for joint reconstruction with implicit denoising to achieve two orders of magnitude faster reconstructions while preserving biologically relevant spatial and temporal structure.

Interpretability analyses revealed that LUCID selectively leverages informative spatial features and angular perspectives, underscoring that the network learns biologically meaningful representations rather than hallucinating structure. Validation experiments across stem cell–derived spheroids and freely moving *C. elegans* demonstrated that LUCID-enabled FLFM provides coherent volumetric reconstructions at sufficient temporal resolution to quantify cell dynamics, tissue strain, and neuronal activity. Furthermore, we extended the framework to bioluminescence imaging, enabling visualization of calcium activity in individual neurons at short exposure times and facilitating volumetric imaging under excitation-free, photon-limited conditions.

Together, these results establish LUCID-enabled FLFM as a versatile and powerful platform for high-speed volumetric imaging under challenging low-SNR conditions. By bridging optical engineering with deep learning, our approach unlocks new opportunities to study fast biological dynamics, from tissue morphogenesis and mechanical strain to calcium signaling, in both fluorescent and bioluminescent systems. This work not only provides a technical foundation for future improvements in light field microscopy but also opens a path toward noninvasive, functional imaging of living systems with subcellular resolution.

## Materials and Methods

### Instrument design

A custom-built optical microscope^5^ was re-equipped with a transmitted LED light source (Thorlabs 590nm), a motorized stage and PiFOC for fast z-acquisition (Piezosystems Jena). Imaging was performed through a 40x 1.25 SiL immersion lens (Olympus # N4274100) and a single-photon resolving qCMOS camera (Hamamatsu Orca Quest, 4.5 um pixel size). An epifluorescence path was added to enable multimodal acquisition of paired bioluminescent/fluorescent images for neuronal network training purposes using a Thorlabs LED. For Fourier light field imaging we redesigned the optical microscope introducing a dual optical branch to switch from a standard widefield microscope to Fourier light field imaging (Fig 2a). For this, we used a 1:1 relay lens with two AC508-150-A-ML lenses with a focal length of *f_F l_*=150 mm to transpose the Fourier plane from the objective into the microlens array (MLA). The selected MLA for our system was the Okotech APH-Q-P1000-R14 with a focal length, pitch, size and radius of curvature of *f_MLA_*=30.6 mm, *P* =1 mm, 14 mm × 14 mm × 1.5 mm and 14 mm respectively, leading to an effective NA per lenslet of *NA_MLA_* 0.016. We chose these specific properties to match the resolution we needed for our particular applications according to

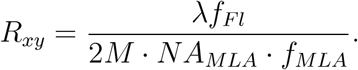

Theoretically, this system, using classical reconstruction methods, is expected to achieve performances of 2.1 *µ*m (*R_xy_*), 16.5 *µm* (*R_z_*), 88 *µm* (DoF), and 147 *µm* (FOV) and an effective magnification of 8.2x. To finish aligning the focal plane of the microlens array with the image plane, we used a relay lens composed of two lenses of 100 mm focal length AC508-100-A-ML. For the widefield light path, we used two right-angle mirror cage (Thorlabs, KCB2EC) with elliptical mirrors and a 200 mm tube lens to generate the image. To switch from one optical branch to the other one, redirecting the light, we employed two motorized flipper mirrors (Thorlabs, 8893-K).

### Sample preparation

Throughout, ‘non-invasive’ denotes the absence of external optical excitation. To enable bioluminescent readouts in *C. elegans*, we employed *bus-17(br2)* and FFz delivery (2% DMSO/0.1% Triton). Bus-17 mutant animals have previously been known to cause skiddy locomotion on agar surfaces due to altered cuticle structure.

#### Caenorhabditis elegans

Animals were maintained using standard protocols^40^ and grown at 20°C. Either larva at the fourth stage (L4) or young adults (YA) with the desired transgene were selected. For fluorescence imaging, worms were transferred to a drop of M9 buffer previously added on top of a 2% agarose pad.^41^ For bioluminescence imaging worms were transferred to a drop of buffer containing 0.2 M citric acid, 0.4 M sodium phosphate dibasic, 2% dimethylsulfoxide (DMSO) and 0.1% Triton X-100 previously placed onto a 2% agarose pad. As substrate for the luciferase, fluorofurimazine (FFz)^42^ 20 mM prepared in PBS was added to the worm suspension in a 1:1 proportion (10 mM final concentration). Transgenic strains are described in Supplementary Table 2. To restrict motion when exposure times increased, neuronal preparations in Fig. 6e–f, animal immobilization on agar pads. We therefore quantified behavioral and calcium baselines under these conditions.

##### Transgenesis. Molecular biology

All plasmids used for this study were constructed using the Gibson assembly method.^43^ Generation of the plasmid expressing the TeNL in glutamatergic neurons and in BWM (pNMSB17 and pNMSB40 respectibvely) have been described before.^44^ Plasmid for expression of the TeNL under the control of *rab-3* promoter (pNMSB133) was constructed by replacing the 15xUAS::delta *pes-10* promoter in pNMSB88 (Addgene plasmid #199321) with the *rab-3* promoter in pHW393 (Addgene plasmid #85583). Primers used for this assembly were 5’-TCCCCGGGATTGGCCAAAGG -3’ / 5’-CCATCGTTGTGAGTGATTTCAGCC - 3’ for pNMSB88 and 5’-TGGCCAATCC-CGGGGATCCTC - 3’ / 5’-GGCTGAAATCACTCACAACGATGGATACG - 3’ for pHW393.

##### Transgenesis

All transgenic animals used for this study were generated by microinjection^45^ into the worm gonad with either of the DNAs previously specified at concentrations 30 ng/µl for pNMSB17, 5 ng/µl for pNMSB40 and 20 ng/µl for pNMSB133. pNMSB17 and pNMSB133 were coinjected with the IR87^46^ as coinjection maker at 5 ng/µl. Plus DNA ladder 1kb from life Sciences up to a final DNA concentration of 100 ng/µl was used as DNA carrier in all cases. For bioluminescence imaging, the transgenic strains were crossed to incorporate the *bus-17(br2)* allele, facilitating the penetration of the substrate required for the reaction.^44^ For the GFP::LMN-1 strain (MSB1136) a split GFP^47^ was performed and the 11th barrel of GFP was introduced by CRISPR/Cas9 at the amino end of the endogenous *lmn-1* gene. The experimental approach was using ribonuclases, repair templates with short homology arms (35bp) and the *dpy-10* system for identification of worms with edits.^48^ Briefly, crRNA in the N-terminal region of *lmn-1* gene (5’ - agaaaaATGTCATCTCGTAA - 3’) was used as well as an ssODN containing left and right homology arms, a flexible linker and the GFP fragment (5’ - tataattaactcttcagaaagcagcgagaaaaATGCGTGACCACATGGTCCTCCACGAGTACGTCAACGCCGCCGGAATCACCGGTGGCGGCAAATTCTCATCTCGTAAAGGTACTCGTAGTTCTCGTATTGTTACGCTAG - 3’). Both components, together with Cas9, tracrRNA were purchased from IDT Technologies. Mix components and concentration in the final mix were: 1.53 µM Cas9, 6.40 µM tracrRNA, 1.25 µM *dpy-10* crRNA, 5 µM *lmn-1* crRNA, 0.92 µM *dpy-10* repair ssODN and 2.20 µM *lmn-1* repair ssODN. The result strain was crossed to CF4588^47^ to reconstitute the full GFP protein.

##### Mouse embryonic stem cells

Stable, bioluminescent cell lines were established as described.^5^ In short, a DNA coding for H2B::TeNL (non calcium sensitive form of the Teal enhanced Nanolantern) was used to integrate into mouse Embryonic Stem Cells. A CRISPR/Cas9 integration into the Rosa26 locus was attempted by the Tissue Engineering Unit at the Center for Genomic Regulation (CRG) in Barcelona. Briefly, CRISPR backbone PX330 was used to introduce the DNA for H2B::TeNL (non calcium sensitive form of the TeNL directed to the nucleus), 5’-ACTGGAGTTGCAGATCACGA - 3’ was used as gRNA and mESC G4, F1 hybrid, superior developmental potential^49^ as host cells. Full insert size was 4516 bp and right and left homology arms were 822 bp and 792 bp long respectively. Transfection was performed on 1 million G4 mESC using Lipofectamine™ 3000 Transfection Reagent (Thermo Fisher Scientific, ref. L3000001) mixed with 2.5 µg of total DNA, 0.625 µg of gRNA and 1.875 µg of circular HDR vector. Lipofection was carried out using a PenStrep-free media. Lipofectamine was removed after 24h. 5 days after lipofection, fluorescence was visible using an EVOS CFP cube. A first round of bulk FACS enrichment was performed. Nozzle 100 µm, To-Pro3 (Thermo Fisher Scientific, ref. T3605) as viability dye. 1.37% of cells were positive for mTurquoise2. 13 days after lipofection, single cells were isolated using FACS and the same conditions as before. The 40% of cells positive for mTurquoise2 were sorted 5×96 plates, resulting in a total of 480 single clones. Screening revealed that 36 out of the 480 clones were expressing fluorescence but PCR testing with primers for the Rosa26 locus 5’ - GGAAAAGTCTCCACCGGACG - 3’ / 5’ - TTGCATTCCAAAAGGAACCACC - 3’ showed that integration had taken place randomly.

mESCs were maintained and expanded on 0.2% gelatin-coated dishes in mESC media composed of the following: Knock-Out DMEM (Thermo Fisher Scientific, 10829-018) supplemented with 15%Fetal Bovine Serum (FBS) (in-house mESC-tested), 1,000 U/ml Leukemia Inhibitory Factor (LIF, in-house generated), 1 mM Sodium Pyruvate (Thermo Fisher Scientific, 11360070), 1x MEM Non-Essential Amino Acids Solution (Thermo Fisher Scientific, 11140050), 50 U/ml penicillin/streptomycin (Thermo Fisher Scientific, 15140-122) and 0.1 mM 2-mercaptoethanol (Thermo Fisher Scientific, 31350010). Cells were cultured at 37°C with 5%CO2. Tissue culture medium was changed every day and cells were passaged using 0.05% Trypsin-EDTA (Thermo Fisher Scientific, 25300054) and quenched 1:5 in DMEM supplemented with 10%FBS (Life Technologies, 10270106). Cells were tested monthly for mycoplasma contamination by PCR.

Spheroids were formed using ultra-low attachment (ULA) round-bottom 96-well plates were used to generate uniform spheroids. Cells were seeded at 5000 cells per well and allowed to aggregate for 24–48 h under standard culture conditions (37*^◦^*C, 5% CO_2_).

Spheroids were then collected and maintained in ES medium for imaging or downstream assays. For differentiation into embryoid bodies, LIF was removed, and spheroids were imaged in the lowlight microscope, in CO_2_ independent medium and at 37*^◦^*C using stagetop incubator (LCIBio).

##### Zebrafish

Zebrafish (Danio rerio) were maintained as previously described.^50^ Embryos were kept in E3 medium at between 25° and 31*^◦^*C before experiments and staged according to morphological criteria^51^ and hours post fertilization (hpf). 4hpf embryos were obtained from the transgenic background with a stable nuclear GFP expression (Tg(actb2:H2B-GFP)).^52^ All protocols used have been approved by the Institutional Animal Care and Use Ethic Committee (PRBB–IACUEC) and implemented according to national and European regulations. All experiments were carried out in accordance with the principles of the 3Rs (replacement, reduction, and refinement).

#### Fluorescence and Bioluminescence imaging

##### Image acquisition of mouse embryonic stem cells (mESCs)

Days prior to imaging, 10^6^ transgenic mESCs (expressing H2B::Teal-enhancedNanolantern, H2B::TeNL) were plated on a 35 mm No. 1 glass bottom *µ*-Dish (Cellvis, D35-10-1-N) coated with 0.1% gelatin (Stem Cell Technologies, 07903). Cells were 40-85% confluent during imaging, providing variation in 3-dimensional colony (spheroid) volume and size. Cells were washed with PBS and new mESC media was added immediately prior imaging to eliminate debris. Imaging was performed at phsyiological conditions using environmental control to maintain 37*^◦^*C and 5% CO_2_ within a top stage incubator (LCI incubator system, Gas Mixer CA-10 and temperature controller TP10). Image acquisition was performed using qCMOS Orca quest C15550-20UP camera with area mode and exposure times depending on the expression strength. Excitation was performed using a 430 nm LED (Thorlabs M430L5) and a standard CFP filter set.

For bioluminescence imaging 2-4 *µ*M fluorofurimazine (FFz, Promega) subtrate was perfused through the cell culture dish, using a peristaltic pump, at a rate of 1 mL/min. Bioluminescence was observed immediately following substrate perfusion. For timelapse imaging, fresh substrate was added every 45-60 mins. Z-stacks of 5 *µ*m spacing were acquired every 5 minutes with an exposure time of 500 ms per slice.

##### Image acquisition of zebrafish embryos

At sphere embryonic stage (4 hpf), embryos stably expressing nuclear GFP (Tg(actb2:H2BGFP)) were mounted in a 35 mm No. 1 glass bottom *µ*-Dish (Cellvis) using 1 % low melting point agarose (Promega) in Danieau’s solution (58 mM NaCl, 0.7 mM KCl, 0.4 mM MgSO4, 0.6 mM Ca(NO3)2 and 5 mM HEPES). Before agarose polymerization, embryos were oriented with blastomere cells facing to the glass. Imaging was performed using a 40×/1.25-NA silicon oil-immersion objective and qCMOS Orca quest C15550-20UP in area mode every 3min for 30 min with 200ms exposure time. Fluorescence excitation was achieved with a Thorlabs LED (M490L4, 490± 26 nm) and a standard GFP filter cube for emission collection.

### Image acquisition of Caenorhabditis elegans

Fluorescence imaging of adult worms was performed using transgenic strains expression LMN-1::GFP as a nuclear signal. Animals were mounted as described^53^ in a 2% agar pad to allow for animal movement and recording of their body-posture dependent subcellular morphology. For immobilized worms, animals were mounted on 6% agar pads. Fluorescence excitation was achieved with a Thorlabs LED (M490L4, 490± 26 nm) and a standard GFP filter cube for emission collection.

For bioluminescence imaging L4 or young adult animals were mounted on a 2% agar pad and treated with citrate buffer (pH 6.5), 20% DMSO, 0.05% triton X-100 and 1.25% pluronic F128 and the chemical cofactor hikarazine^54^ for the generation of a bioluminescence signal. Bioluminescent signals from neuronal and body-wall muscle activity were recorded from young adult animals were mounted on a 2% agar pad, and 1 *µ*L of 20 *µ*M fluorofurimazine (FFz, Promega) was applied immediately before imaging, without any preincubation.

### Richardson Lucy Deconvolution for image reconstruction

Volumetric reconstruction of Fourier Light Field Microscopy (FLFM) data was performed using a model-based Richardson-Lucy (RL) deconvolution framework, adapted from the computational methods previously described.^7,12^ The reconstruction algorithm uses a forward model H representing the FLFM imaging system as a spatially varying convolution over axial depth, with each depth plane encoded by a 2D point spread function (PSF). The PSF stack *H* and its adjoint *H*^T^, either experimentally measured or synthetically simulated based on the optical geometry, were preloaded into memory as depth-resolved kernel banks.

For a given raw lenslet image stack *I*, the forward model simulates the measured image by convolving each axial slice of an estimated object volume *V* with its corresponding PSF, and summing over depth. The inverse problem is solved iteratively using the RL update rule:

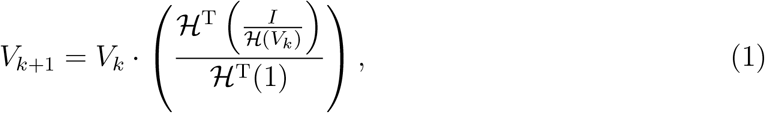

where H and H^T^ denote forward and backward projection operators, implemented through FFT-based convolutions on a GPU to accelerate computation. The object volume was initialized with a uniform guess and iteratively refined over 30 RL iterations. To minimize edge artifacts and reduce the data footprint, a centered sub-volume of fixed dimensions was cropped and saved every 10 iterations. The reconstruction yielded a 3D estimate of the object.

### Attention neural networks for the reconstruction of Fourier light field images

We developed a deep learning-based pipeline for real-time volumetric bioluminescent imaging using an attention-based convolutional neural network (CNN) trained on synthetic, fluorescent and bioluminescent data. To generate the training dataset, we designed a dual-branch optical system that provides corresponding 3D wide-field images of immobilized samples, capturing their Fourier view perspectives using a micro-lens array (MLA). The images are acquired sequentially by switching between branches using motorized flip mirrors (Fig. 2a). For training, each perspective was used to extract features, which were then integrated into subsequent convolutional layers. The network incorporates two attention mechanisms to preserve high-frequency information for enhanced reconstruction quality (Fig. 2b). First, a view-attention branch is added to each encoder block, addressing the loss of high-frequency content due to max-pooling layers. Second, attention gates are applied in the skip connections to combine spatial and angular information from the down-sampling and up-sampling paths. These soft-attention mechanisms reduce the activation of irrelevant input regions, minimizing redundant features.^29^ The final layer of the network is configured to produce a specific number of slices that match the depth of the 3D target stack. We used a weighted loss function that combines two terms: pixel-wise mean squared error (MSE) and edge-aware error. The edge-aware term is computed by applying the Sobel operator to both the reconstructed and reference images and then taking the MSE between their edge maps. This encourages the network to preserve high-frequency structures, rather than simply weighting pixel errors near edges.^11^ This combination drives the network to not only infer accurate intensity distributions but also generate sharper structures, resulting in higher resolution reconstructions.

### Synthetic forward model

To generate synthetic training pairs, we treat the experimentally acquired wide-field volume as a reference volume, *V*_ref_ , and project it through the calibrated Fourier light-field forward operator, H_FLFM_, to obtain the corresponding synthetic light-field input:

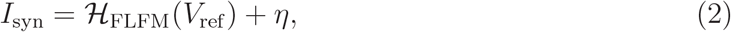

where *η* denotes measurement noise. Importantly, *V*_ref_ is a WF-derived experimental reference volume and is not treated as an absolute ground truth volume.

### Loss function

The network was trained with a weighted volumetric loss consisting of a voxel-wise reconstruction term and an auxiliary edge-preserving regularizer:

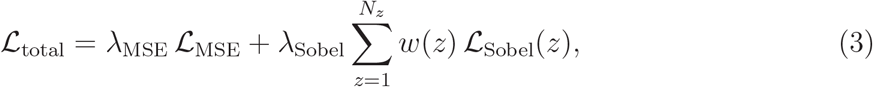

where L_MSE_ is the primary reconstruction loss, L_Sobel_ is the Sobel-based gradient loss computed slice-wise, and *w*(*z*) is a Gaussian weighting function centered on the focal plane:

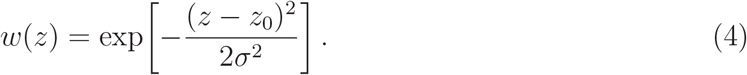

Here, *z*_0_ denotes the focal plane and *σ* controls the axial extent of the weighting.

### Training procedure for Fourier light field data

In the case of synthetic Fourier light field data, we first collected N different 3D stacks (of size 4096 px × 2304 px × 101 planes). The acquisition of the 3D stacks was made using an epifluorescence microscope with a 40×/1.25-NA silicon oil-immersion objective and a 430 nm excitation wavelength (Supplementary Figure 2a). We cropped the ROI from the acquired stack to account for the iris positioned in the image plane, which limits the FOV. As a result, all obstructed areas were removed to optimize computational memory usage. The same as before, we augment the synthetic data by performing linear transformations, e.g. flipping, rotating, and inverting the z axis to augment the dataset. We calculated the point spread function of the synthetic Fourier light field (PSF) based on wave optics theory by performing a forward projection onto the high resolution stacks^20^ to generate the corresponding synthetic Fourier light field image to train the network. The synthetic PSF was calculated to match our experimental setup; in other words, the synthetic PSF had the same amount of elemental images (EIs) as our experimental setup. The network training process follows a methodology similar to our earlier work,^5^ where extracted views from the original light field image are transformed into a conventional 3D stack. We trained the network with a patience parameter of 10 epochs, which means that the training stopped if the loss function did not improve over 10 consecutive epochs. This training process required approximately 5 hours on a single graphical processing unit. As before, reconstruction performance was assessed using structural similarity (SSIM) and normalized root means squared error (NRMSE) metrics on unseen data.^5^

An initial model was trained on synthetic Fourier light field datasets and subsequently refined through transfer learning using experimentally acquired FLFM data. This procedure enables the network to adapt to the noise characteristics, photon statistics, and optical aberrations present in experimental images, improving reconstruction accuracy and reducing artifacts associated with synthetic-only training.

The network training and data preparation consisted of the following steps:

- **Synthetic PSF generation:** A synthetic point spread function (PSF) was computed to match the experimental configuration.^20^ Specifically, the number of elemental images (EIs) in the synthetic PSF matched the number in the experimental FLFM dataset.
- **Initial network training:** Following the approach of our previous work, ^5^ perspective views were extracted from raw Fourier light field (FLF) images and restructured into conventional 3D stacks. The model was trained with a patience parameter of 10 epochs, terminating training if the validation loss did not improve for 10 consecutive epochs.
- **Transfer learning with experimental data:** Since a model trained solely on synthetic data introduced artifacts, we refined it using experimental data through transfer learning. Using the FLFM arm of the low light scope, we acquired a fluorescent image of *C. elegans* after direct excitation fo the mTurquoise fluorescence protein of the TeNL luciferase under the control of the pan-neuronal *rab-3* promoter. The corresponding high-resolution 3D reference stack was obtained with the widefield (WF) imaging arm on the same sample.
- **Perspective view extraction and calibration:** Raw FLFM images were processed to extract angular views. This required subpixel-accurate calibration of microlens positions, achieved by illuminating the sample with a homogeneous light source. A custom routine then automatically identified the lenslet array and extracted angular views from the raw image.
- **Preprocessing and augmentation:** The extracted perspective views were aligned and prepared as input–output pairs for supervised learning. We applied data normalization and augmentation to enhance generalization and reduce overfitting. The first stage of the network included a subpixel convolutional upsampling layer to enhance resolution while extracting relevant image features.
- **Widefield stack preparation:** WF images were cropped due to the pinhole located at the center of the relay lens, keeping only regions with relevant signal. The original WF volume (1300×1300×101 voxels) was downsampled to 800×800×101 to reduce computational burden without sacrificing structural integrity. This was further adjusted to 200×200×101 for compatibility with the model’s transposed convolutional layers, allowing for a precise 4× up-sampling.

Neural networks were implemented in TensorFlow/Keras (v2.12) and trained using the Adam optimizer with a learning rate of 1e-4 and a batch size of 32. Training was performed for up to 250 epochs with early stopping (patience = 10 epochs), terminating when the validation loss did not improve for 10 consecutive epochs. The validation set comprised 10% of the training data and was randomly selected without overlap with either the training or test sets. Dataset partitions were chosen to capture the variability observed across experimental time points. All training procedures were repeated at least three times to ensure reproducibility. Training was performed on a workstation equipped with an Intel Xeon Gold 6248R CPU, 128 GB RAM, and an NVIDIA Quadro RTX8000 GPU (48 GB memory), requiring approximately 5 hours per model.

We note that wide-field (WF) stacks are used as experimental reference volumes rather than absolute physical ground truth. Due to out-of-focus contributions inherent to diffraction-limited WF imaging, these references do not represent physical volumetric truth. Consequently, similarity metrics (e.g., SSIM) should be interpreted as agreement with the reference modality rather than absolute accuracy representing the physical object.

### 3D reconstruction with bioluminescence Fourier light field data via transfer learning

Volumetric reconstruction of bioluminescent Fourier light field (FLFM) images presents additional challenges compared to fluorescence imaging due to significantly lower photon counts and reduced signal-to-noise ratio (SNR). To address this, we generated an experimental bioluminescent dataset using *C. elegans* expressing mTurquoise2 pan-neurally with the *rab-3* promoter, acquired at varying exposure times (1 s and 5 s). These FLF images - characterized by inherently lower SNR - were paired with corresponding widefield (WF) stacks serving as reference for supervised training (Fig. 6).

Initial models pre-trained on fluorescence data were insufficient for bioluminescence, often producing blurry or noisy reconstructions that lack fine structural detail. To improve performance, we used a three-stage training strategy: (1) pre-training on synthetic data, (2) fine-tuning on fluorescence experimental data, and (3) final training on bioluminescence experimental data. This approach enabled the model to learn denoising and reconstruction jointly, resulting in markedly improved volumetric fidelity (Fig. 6). To perform the transfer

learning, we loaded the weights of the pretrained network with synthetic images and train it again with the experimental dataset using the same hyperparameters stated before (Fig. 2). In contrast, models trained solely on synthetic, fluorescence, or bioluminescence data performed poorly, frequently missing key features or amplifying noise (Fig. 6). Reconstruction quality was subsequently evaluated with respect to structural fidelity, noise suppression, and processing time and physiological calcium dynamics (Fig. 6).

### Calculation of the Signal/Noise ratio (SNR)

To prove the improvement of the SNR after the restoration with the neural networks, we calculated the SNR of the input and the processed image. Here, we define the SNR as the ratio of the mean of the background subtracted signal intensity over the standard deviation of the background noise,

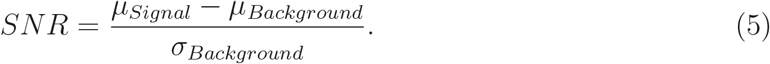

To select what was considered as noise and what was considered as signal, we sorted the intensities present in the image and calculate the gradient of the list of intensities. The cutting point was selected where a high inflection gradient was found in the distribution of intensities. The SNR computation only considered the pixels above that inflection point as signal and the rest as background.

### Calculation of the spatial frequencies

Spatial-frequency content and effective image resolution were assessed using the parameter-free decorrelation analysis described by Descloux et al.^21^ The method analyzes partial phase correlations in Fourier space to identify the highest spatial frequency, *k_c_*, containing sufficient signal relative to noise. The corresponding effective resolution was calculated from *k_c_* according to *R* = 2*p/k_c_*, where *p* is the image pixel size. Decorrelation curves and the derived resolution estimates were compared between WF reference images and LUCID reconstructions to assess preservation of spatial-frequency information.

### Associated content

#### Supplementary material is available for this manuscript

Additional experimental details; optical calibration and microlens-registration procedures; reconstruction parameters; training/validation settings and hyperparameters; Supplementary Figures S1–S9; Supplementary Tables S1–S3; and Supplementary Videos S1–S8 (AVI, MP4)

#### Data and Code availabilty

The data supporting the findings of this study are available within the Article and its Supporting Information. Raw image data, trained model weights, and versioned reconstruction code are openly available at https://gitlab.icfo.net/NMSB/lucid. We deposited the design of the FLFM under 10.5281/zenodo.17199808

#### Notes

The authors declare no competing financial interest.

## Acknowledgement

The authors thank Aleksandra Pidde and Diego Ramallo for advice on programming, Gustavo Castro and Pablo Loza for help with the optical design and Montserrat Porta-de-la-Riva, Santiago Ortiz and Malak ElQuessny for help with biological sample preparation (*C. elegans*, fish and mESCs, respectively). We thank Verena Ruprecht lab at the CRG Barcelona for sharing zebrafish embryos for Figure 1 d-f and Santiago Ortiz for help with imaging.

MK acknowledges financial support from the ERC (PoC ”LowLiteScope”, 101138041), Human Frontiers Science Program (RGP021/2023), MCIN/ AEI/10.13039/501100011033/ FEDER “A way to make Europe” (PID2024-157334OB-I00), “Severo Ochoa” program for Centres of Excellence in R&D (CEX2019-000910-S), from Fundació Privada Cellex, Fundació Mir-Puig, and from Generalitat de Catalunya through the CERCA and Research program and funding through H2020 Marie Sk-lodowska-Curie Actions (847517 to LFMC). ICFO is the recipient of the Severo Ochoa Award of Excellence of MINECO (Government of Spain). This work was supported in part by Oracle Cloud credits and related resources provided by Oracle.

## For Table of Contents Use Only

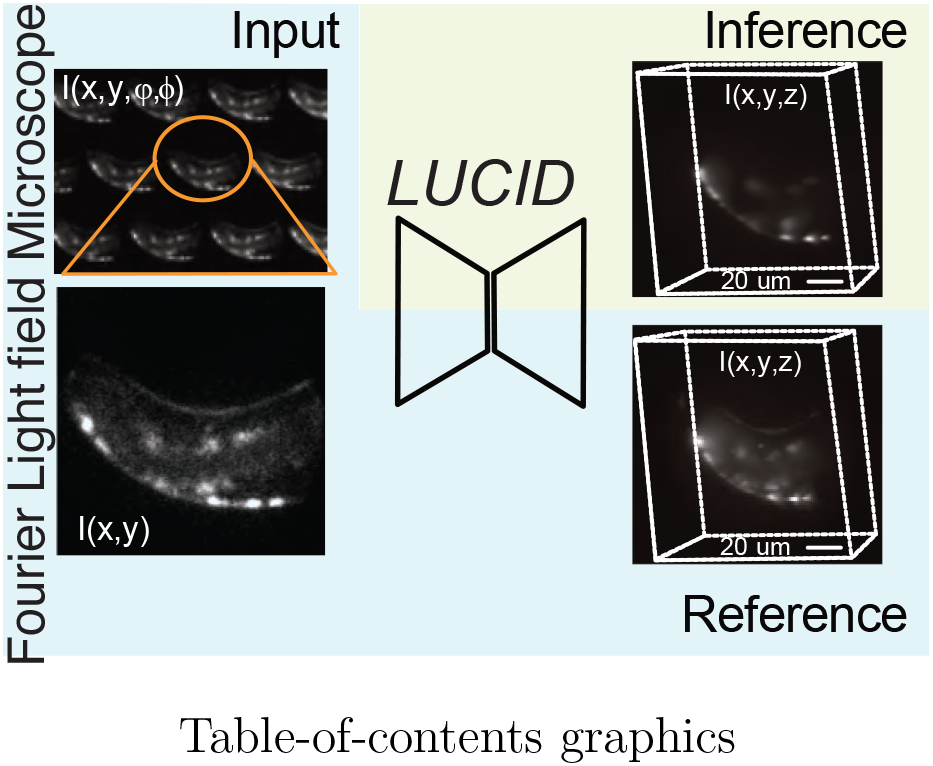

## Supporting Information Available

### Supplementary Figures

**Supplementary Fig. 1:**
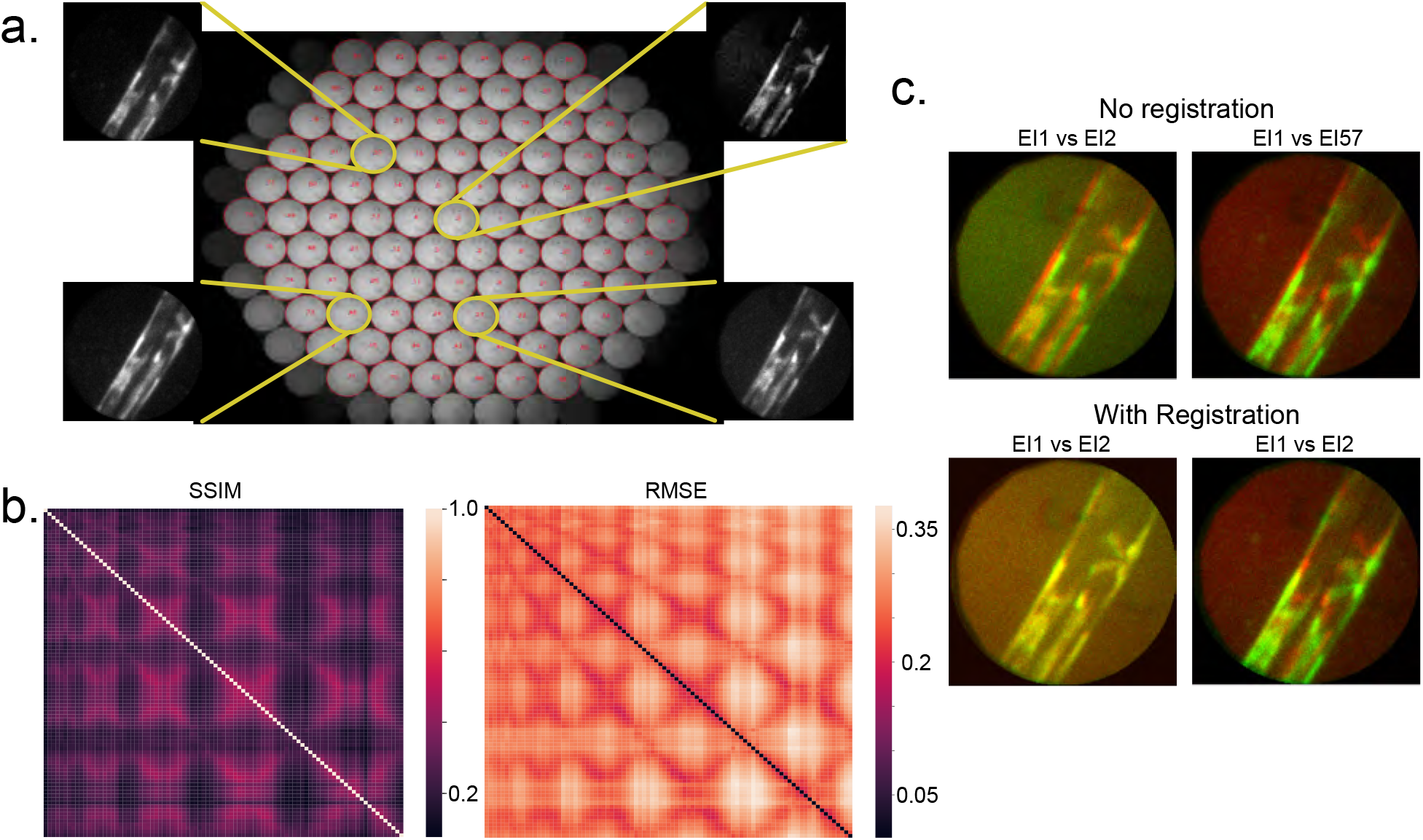
Analysis of perspectives in FLFM. **a**. Lenslet distribution showcasing four representative elemental images (numbered 0, 23, 30, and 46), each capturing unique information of *C. elegans* body wall muscle. **b**. SSIM and RMSE heatmaps, highlighting low similarity across most perspectives except for self-comparisons along the main diagonal. **c**. Comparison of elemental images (body wall muscle of *C. elegans*) after rigid registration, ensuring fair evaluation of information uniqueness.

**Supplementary Fig. 2:**
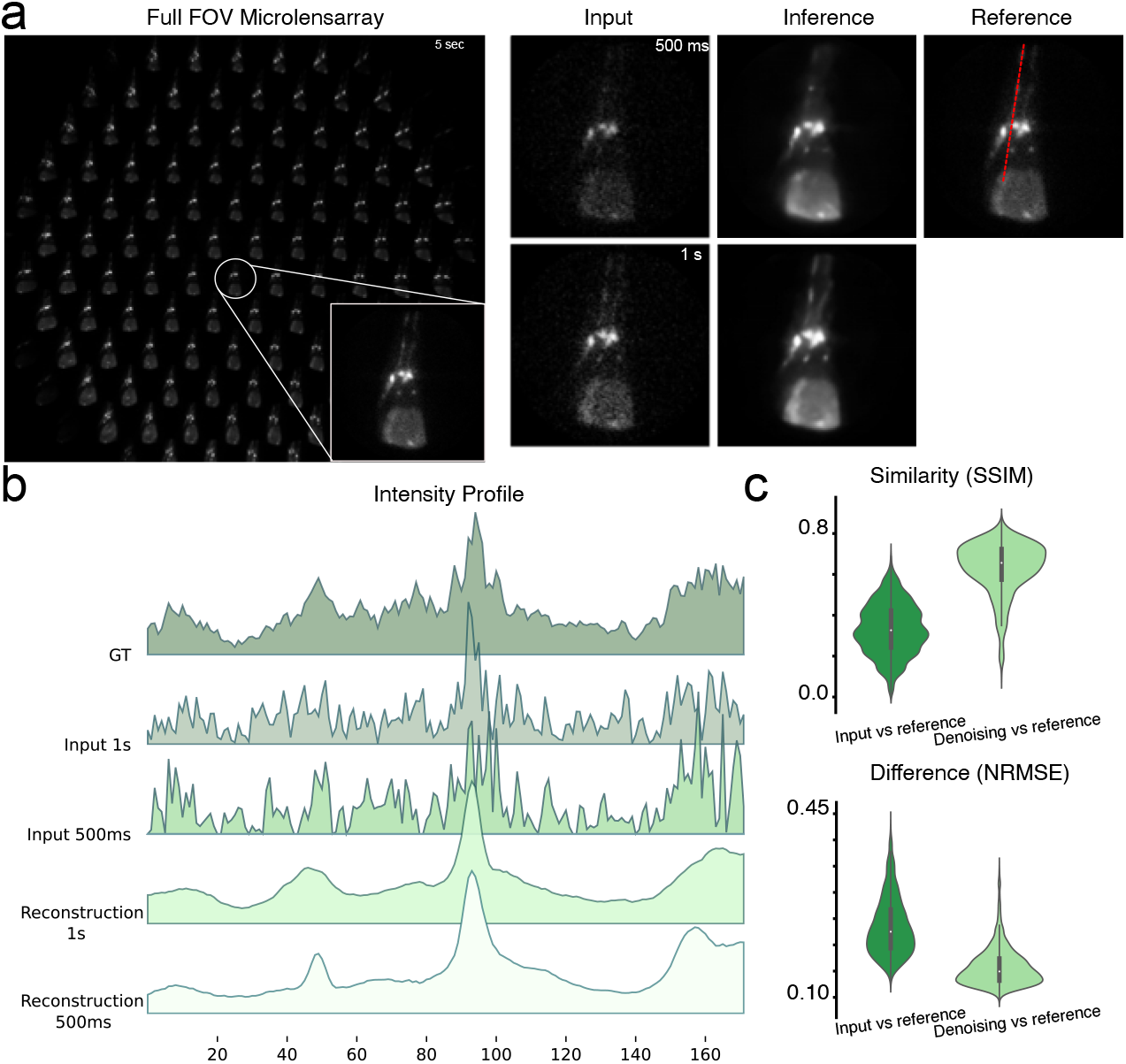
Denoising of Fourier Light Field Elemental Images. **a.** Experimental raw LF image with 91 perspectives of the head of a worm expressing the mTurquoise2 in glutamathergic neurons. Each of the images shows a different perspective from the sample. Noisy image input with exposure times of 500ms and 1s (top and bottom), then their respective restorations using a denoising model and an elemental image taken with a 5s exposure time. Along the head of the animal a red dotted line is shown **b**. to see the intensity profile along the image. The two input images are highly noisy prior to any post-processing. Following restoration, the signal becomes smooth while preserving salient image features, with background noise and other non-informative components effectively suppressed. **c**. The restoration achieves an approximately 2 times of improvement using the SSIM, NRMSE against the reference images. N = 38.

**Supplementary Fig. 3:**
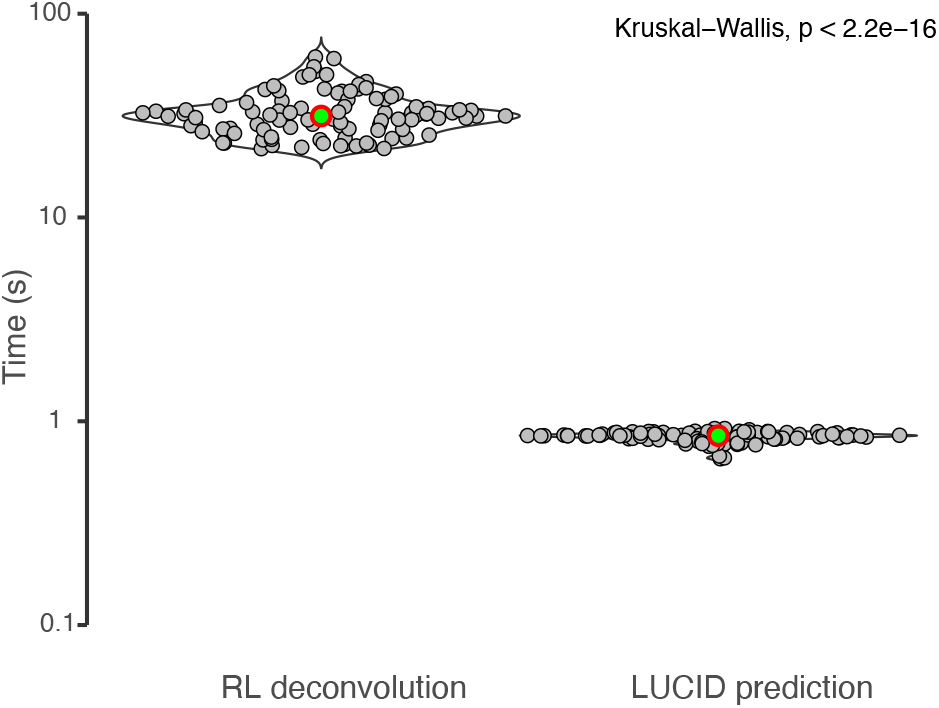
Distribution of Execution Times: Deconvolution vs Neural Network. Violin plot of the duration of the reconstruction of the FLF images using Richardson Lucy and LUCID prediction pipeline.

**Supplementary Fig. 4:**
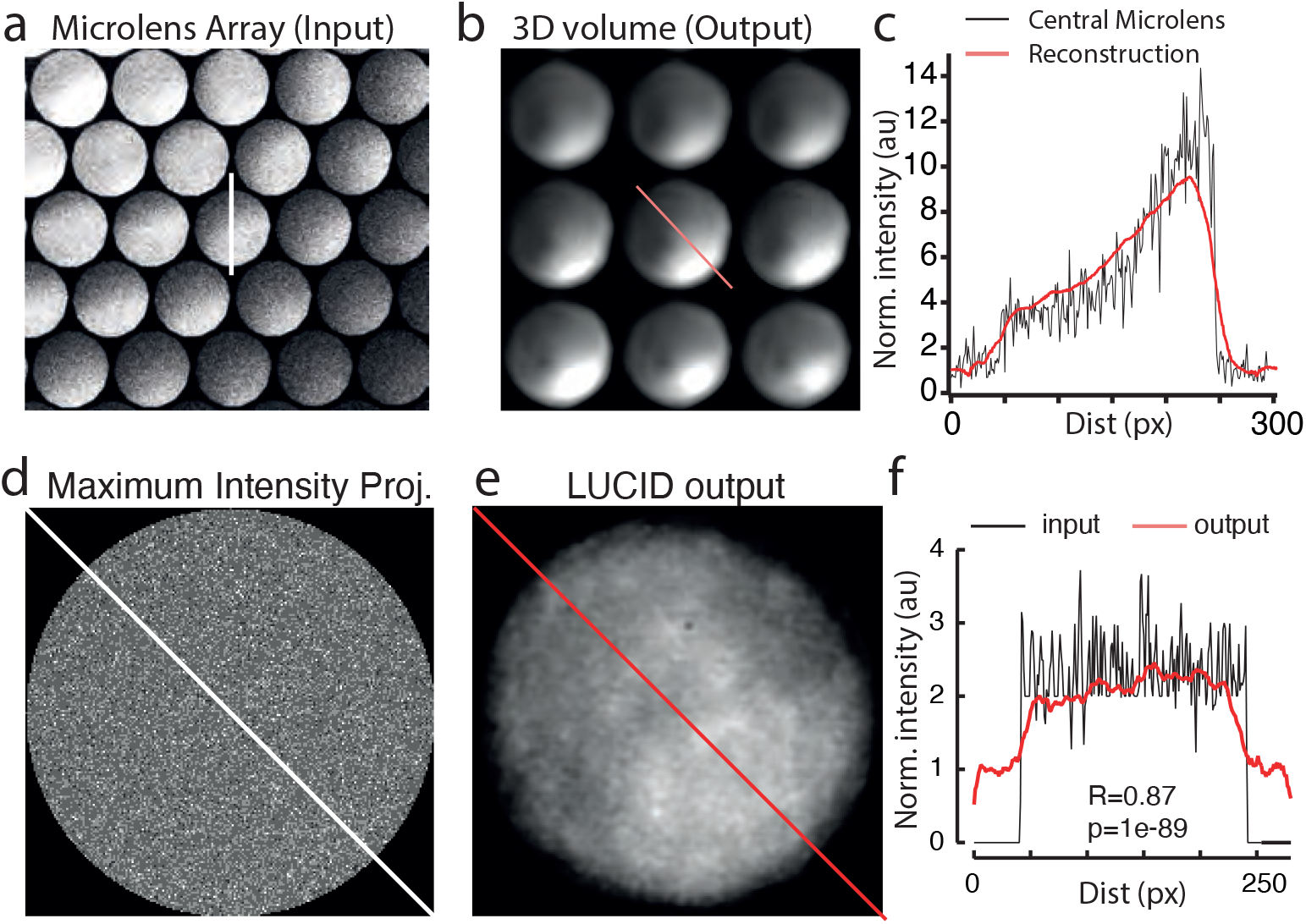
LUCID does not introduce artificial gradients or steps. **a.** Representative image of a sample showing a fluorescent gradient captured with the microlens array. **b.** Reconstruction: Consecutive z-planes at 10 µm intervals from a reconstructed 100 µm volumetric stack from the FLFM image in (a). **c.** Intensity profile along the lines indicated in (a) and (b). **d.** Maximum intensity projection of all elemental images acquired on an unlabeled sample with 200ms exposure time to capture pure camera noise. **e.** Reconstruction: Maximum intensity projection of the reconstructed FLFM image in (a). **f.** Intensity profile along the lines indicated in (d) and (e). R indicates the Pearson correlation coefficient with p as the two-sided p value.

**Supplementary Fig. 5:**
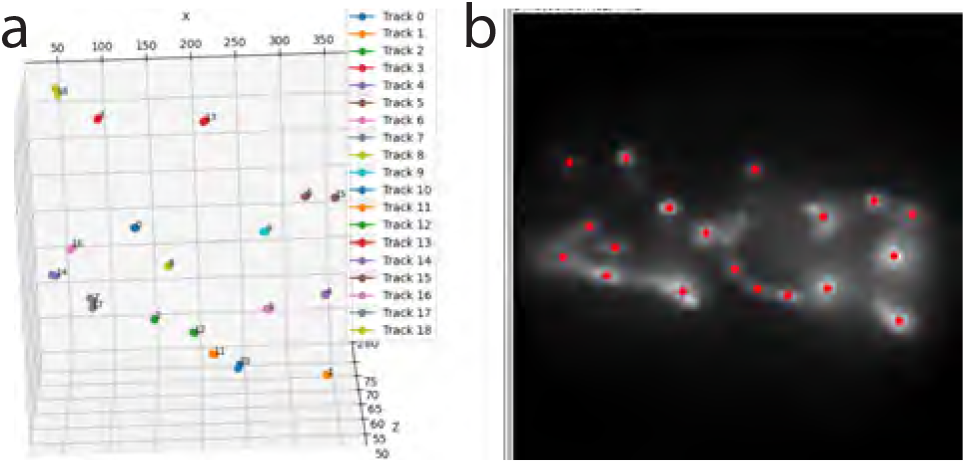
Quantifying nucleus position in a Fourier light field time lapse of representative anesthetized animal. **a,** Graph showing position of the segmented nuclei for all frames of the reconstructed timelapse **b,** Representative maximum intensity projection of a reconstructed volume at a given time point in the movie.

**Supplementary Fig. 6:**
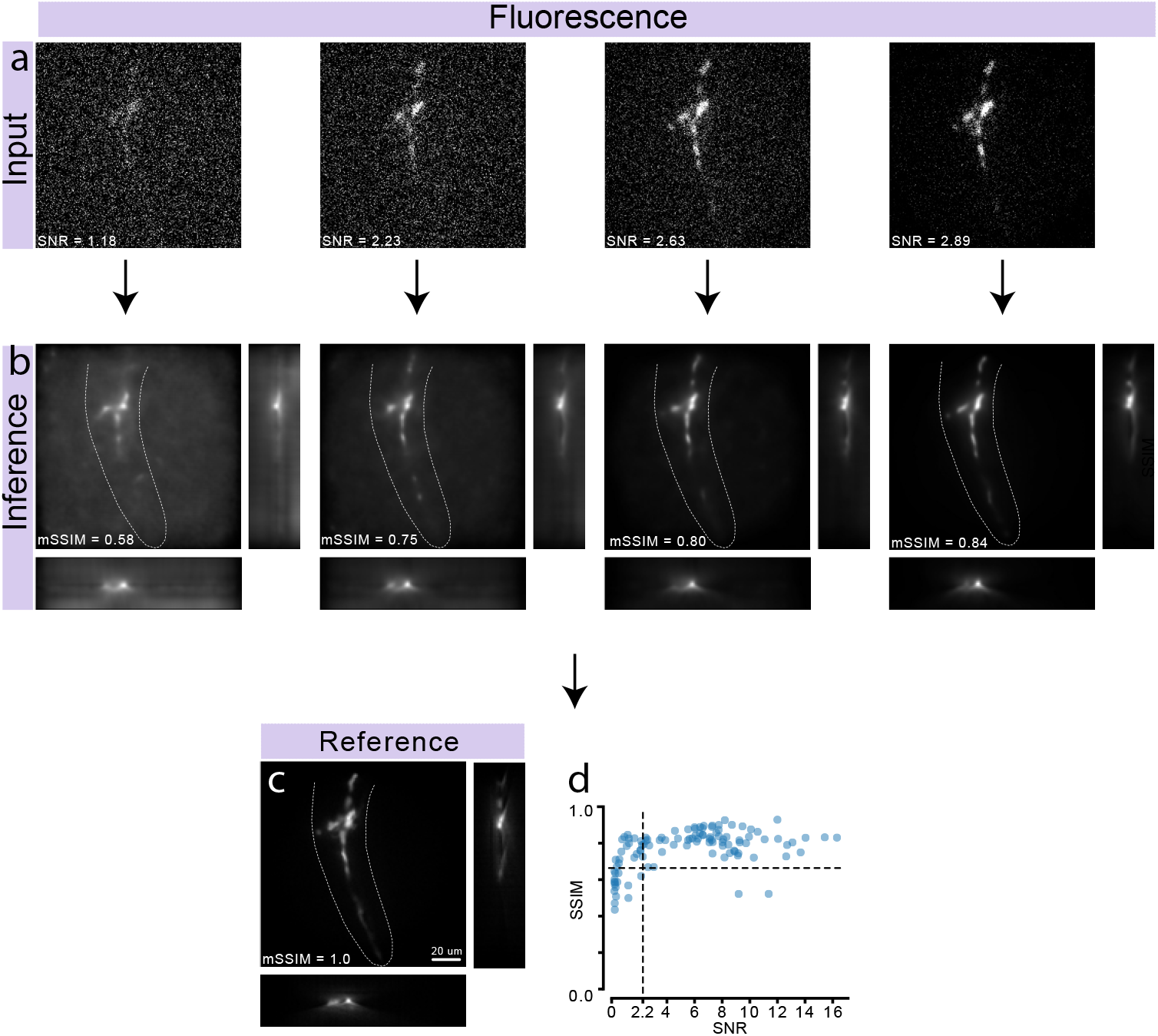
Limits of restoration quality in FLF reconstruction. **a.** Representative input images showing the central EI of fluorescent *C. elegans* animal with neuronal labeling, acquired at different exposure time yielding varying signal/noise levels. **b.** LUCID reconstructions with their respective SSIM compared to the reference. Higher input SNR yields better structural similarity (SSIM) in the restored images. **c.** Reference images studied in **a** and **b**. **d.** SSIM plotted against input SNR, illustrating a threshold for effective reconstruction. Each dot represents a pair of images in the pipeline. SNR = signal-to-noise ratio. Scale bar = 20 *µ*m.

**Supplementary Fig. 7:**
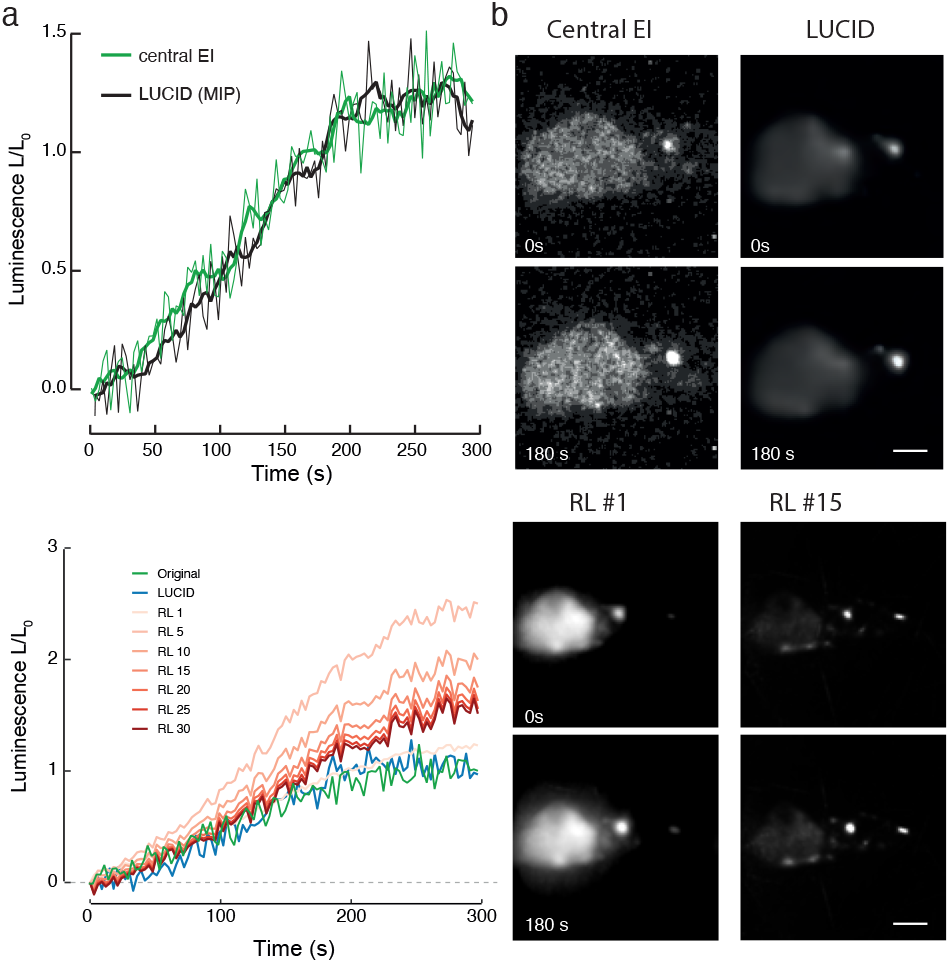
Comparison of Calcium signals between the widefield reference and Lucid reconstructions. **a,** Graph showing the temporal bioluminescence dynamics of the genetically encoded calcium indicator (TeNL) in the DVA neuron, extracted from the central microlens (elemental) images of the MLA and from the LUCID reconstruction. Note, the central elemental view is an internal comparator, not an absolute reference measurement. The thick lines are smoothed using a 10 point median filter. **b,** Representative images from the central microlens and the maximum intensity projection of the reconstruction at the indicated timepoints. Scalebar = 5µm.

**Supplementary Fig. 8:**
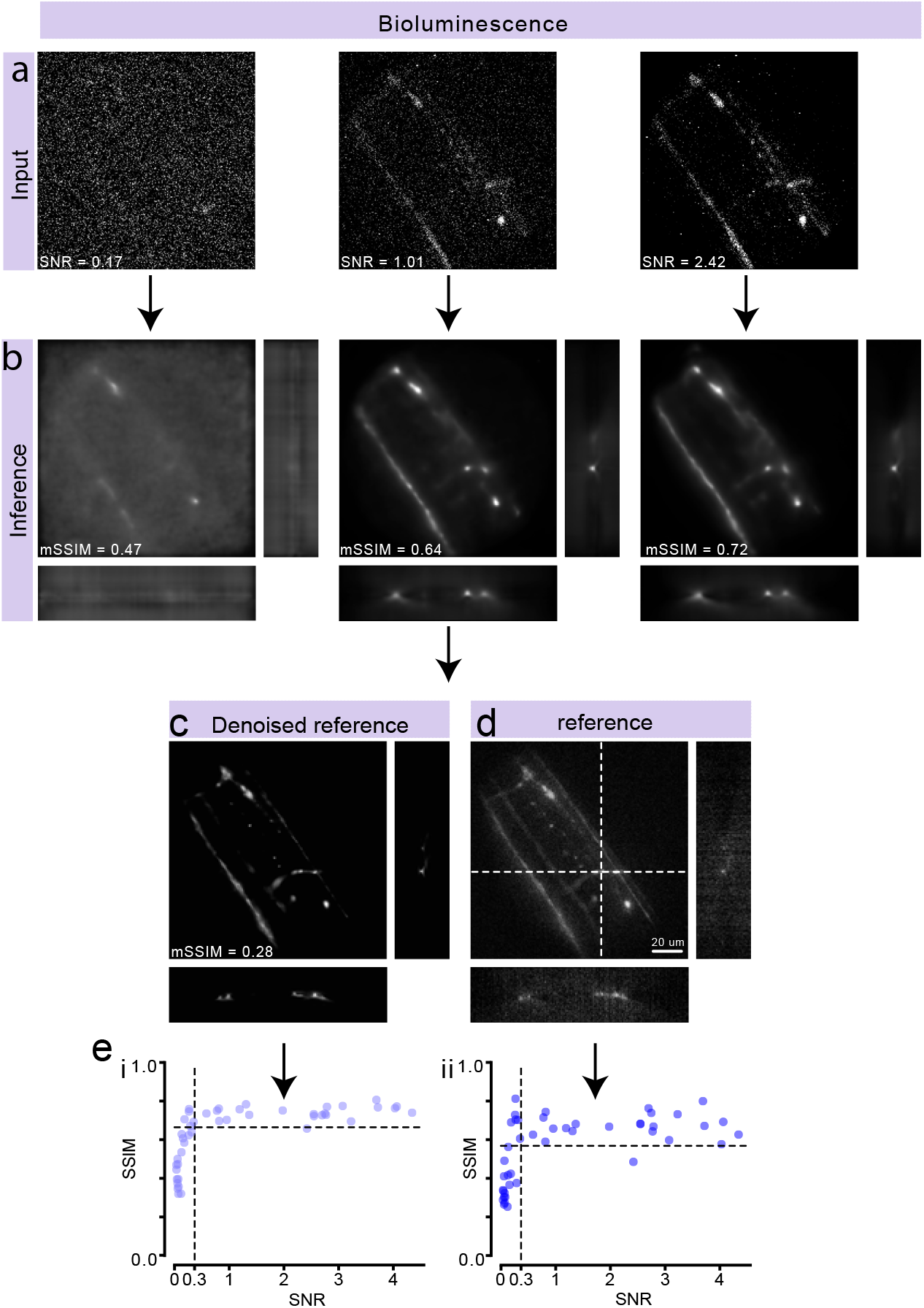
Limits of restoration quality for bioluminescent FLFM reconstruction. **a.** Input elemental images used for reconstruction, corresponding to different SNR levels, illustrating how input noise affects model performance. **b.** Orthogonal views of the FLF reconstructions, displayed alongside their respective mSSIM values. **c.** Orthogonal view of the denoised reference volume, obtained by applying a neural network-based denoising method to the raw reference images, serving as a high-quality reference for comparison. **d.** Orthogonal view of the raw reference volume prior to denoising, showing the noise characteristics present in the original data. **e.** Plot of mean SSIM (mSSIM) versus input SNR, highlighting how reconstruction quality compares when using either the (i) denoised or (ii) raw images as a reference. SNR = signal-to-noise ratio. Scale bar = 20 *µ*m.

### Supplementary Tables

**Supplementary Table 1.** Hyperparameters. Table 1: Summary of the different hyperparameters tested to train the LUCID base model. Overview of the deep learning model configurations assessed in this study for fluorescence FLFM. Each entry includes the number of training images, presence or absence of data augmentation, indication of experimental status, use of transfer learning, network depth, whether any layers were frozen during training, and application of denoising methods employed.

| Model | N im-<br>ages | Augmen-<br>tation | Experi-<br>mental | Transfer<br>learning | Freezing | De-<br>noising | Depth |
| --- | --- | --- | --- | --- | --- | --- | --- |
| 1 | 748 | Yes | Yes | No | No | No | 2 |
| 2 | 748 | N/A | Yes | No | No | No | 3 |
| 3 | 748 | N/A | Yes | No | No | No | 4 |
| 4 | 748 | N/A | Yes | No | No | No | 2 |
| 5 | 1468 | N/A | Yes | No | No | No | 2 |
| 6 | 1468 | N/A | Yes | No | No | No | 4 |
| 7 | 1468 | N/A | Yes | No | No | No | 4 |
| 8 | 2188 | N/A | Yes | No | No | No | 4 |
| 9 | 2188 | N/A | Yes | No | No | No | 4 |
| 10 | 2188 | N/A | Yes | No | No | No | 4 |
| 11 | 2188 | N/A | Yes | No | Yes | Yes | 4 |
| 12 | 2188 | N/A | Yes | Yes | Yes | Yes | 4 |
| 13 | 2188 | N/A | Yes | No | No | No | 4 |
| 14 | 2188 | N/A | Yes | Yes | No | No | 4 |
| 15 | 2188 | N/A | Yes | Yes | No | No | 4 |
| 16 | 2188 | N/A | Yes | Yes | No | No | 4 |

**Table 2.**
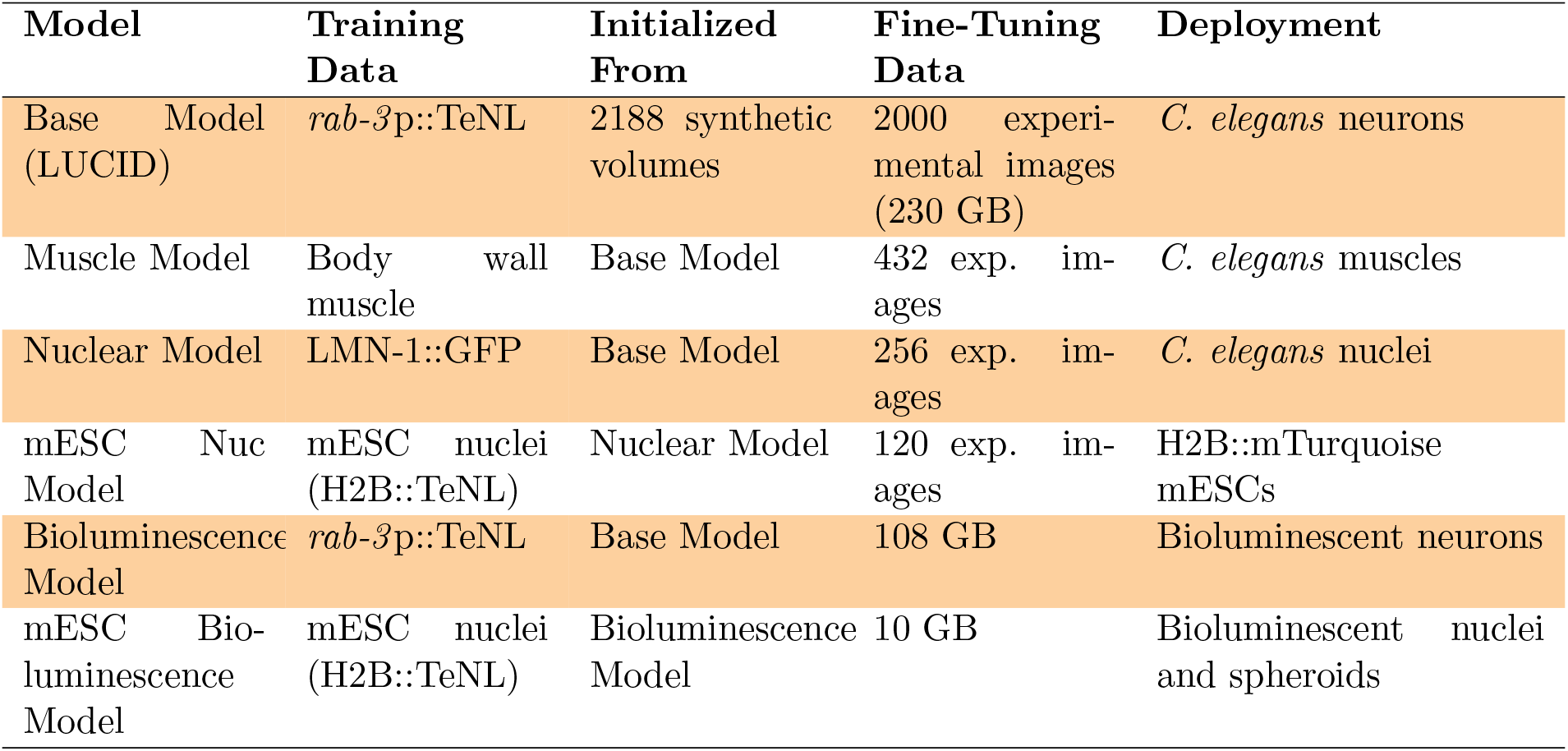
Summary of LUCID models developed in this work through transfer learning.

**Supplementary Table 2.** Composition of the data sets used for transfer learning.

**Supplementary Table 3.**
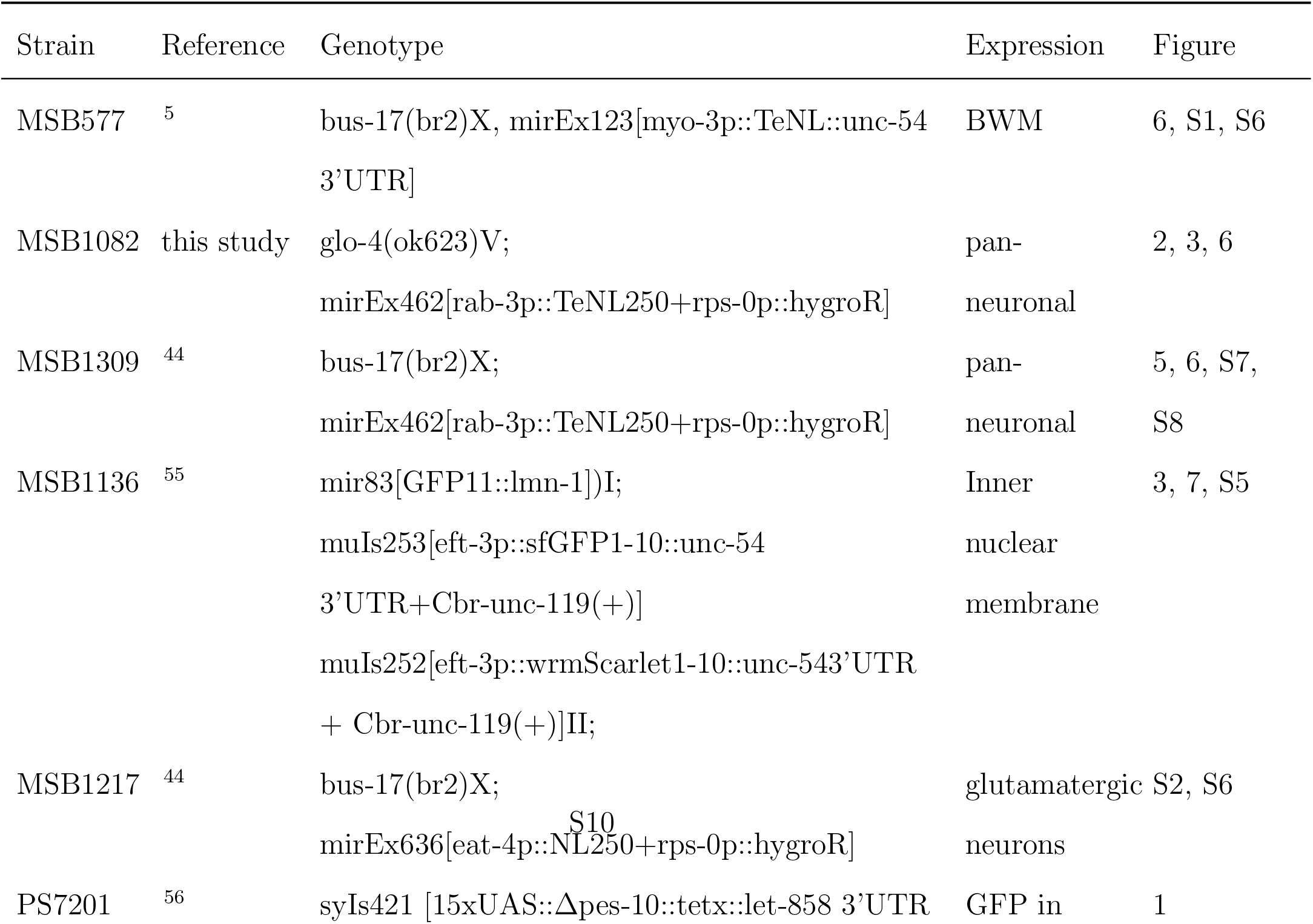
*C. elegans* strains used in this study.

#### Supplementary videos

**Supplementary Video S1**

3D restoration of a short timelapse video of a developing zebrafish embryos at 50% epiboly. Zebrafish are transgenic for histone::GFP, and videos were deconvolved using Richardson-Lucy algorithm. Scale bar = 20 µm. Interframe interval = 3 min. Exposure time = 200ms.

**Supplementary Video S2**

Central elemental image of a freely behaving *Caenorhabditis elegans* animal, transgenic for H2B::Nanolantern under fluorescence contrast.

**Supplementary Video S3**

Short timelapse video of freely behaving *Caenorhabditis elegans* transgenic for H2B::Nanolantern under fluorescence contrast, shown in Suppl. Video S2. Reconstructed using LUCID. Scalebar 20 µm.

**Supplementary Video S4**

Short timelapse video of fluorescently labeled mouse embryo stem cells transgenic for H2B::Nanolantern. Interframe interval = 1 min.

**Supplementary Video S5**

3D restoration of a short timelapse video of freely behaving *Caenorhabditis elegans* transgenic for calcium sensitive Nanolantern in body wall muscles (*myo-3* p::TeNL) under bioluminescence contrast. Restoration was performed with the LUCID model. Scale bar = 20 µm.

**Supplementary Video S6**

A short timelapse video from the central elemental microlens depicting the motorneurons in the ventral nerve chord freely behaving *Caenorhabditis elegans* transgenic for calcium sensitive Nanolantern under the control of the *rab-3* promoter in bioluminescence contrast.

**Supplementary Video S7**

Restoration of a timelapse video of freely behaving *Caenorhabditis elegans* transgenic for calcium sensitive Nanolantern under the control of the *rab-3* promoter in bioluminescence contrast, showing a section of the ventral nerve chord.

**Supplementary Video S8**

Restoration of a short timelapse video of the dorsorectal ganglion in bioluminescence contrast, using a calcium sensitive Nanolantern under the control of the *rab-3* promoter. Anterior signal is the off-target expression of the Nanolantern in the posterior gut. Interframe interval = 3s. Length of the arrow = 40 µm.

## Notes

### Competing Interest Statement

The authors have declared no competing interest.

### Summary of Updates

Refined conceptual presentation, algorithms and analyses.

